# Sampling globally and locally correct RNA 3D structures using ERNWIN, SPQR and experimental SAXS data

**DOI:** 10.1101/2022.07.02.498583

**Authors:** Bernhard C. Thiel, Giovanni Bussi, Simón Poblete, Ivo L. Hofacker

## Abstract

The determination of the three-dimensional structure of large RNA macromolecules in solution is a challenging task that often requires the use of several experimental and computational techniques. Small-angle X-ray spectroscopy can provide insight into some geometrical properties of the probed molecule, but this data must be properly interpreted in order to generate a three-dimensional model. Here, we propose a multiscale pipeline which introduces SAXS data into modelling the global shape of RNA in solution, which can be hierarchically refined until reaching atomistic precision in explicit solvent. The low-resolution helix model (ERNWIN) deals with the exploration of the huge conformational space making use of the SAXS data, while a nucleotide-level model (SPQR) removes clashes and disentangles the proposed structures, leading the structure to an all-atom representation in explicit solvent. We apply the procedure on five different structures up to 126 nucleotides with promising results.

## Introduction

The incorporation of experimental data into the structural modelling of nucleic acids is an increasingly common approach, which helps to scale down the sea of possible structures that computer algorithms can propose. ^1^ Among these experimental techniques, small-angle X-ray scattering (SAXS) is a handy tool for providing insight into the global shape and size of macromolecules in solution,^2,3^ which can be employed to refine models or to guide molecular dynamics (MD) simulations. The information provided by SAXS spectra is, nevertheless, a 1-dimensional intensity profile of limited resolution, which opens the questions of how to interpret and integrate this data into an all-atom model. Although in principle the scattering intensities can be calculated exactly from the atomic coordinates, the effect of the hydration shell is not known in a closed form, and it is usually taken into account through different approximations along diverse tools. For example, an implicit water model is used in PLUMED^4^ and in the original version of CRYSOL, ^5^ but later versions of this software employ dummy water beads to model the hydration shell, ^6^ in the same spirit as Fast-SAXS-pro.^7^ Ahead in complexity, methods such as WAXSiS ^8^ and Capriqorn ^9^ take into explicit consideration the solvent effects, by analyzing the SAXS contribution of an entire set of MD-generated trajectories, and subtracting the solvent effects obtained from an independent simulation of the buffer. In all these approaches, there is a trade-off between speed and accuracy, ^10^ which has to be taken into account when dealing with large RNA structures.

There is also a variety of ways for building molecular models based on SAXS profiles. A common approach is to generate a huge number of structures and to afterwards select those with the best match between the predicted SAXS profile and the experimental data.^11^ SAXS intensities have been also used to reweight or sample from MD simulations. ^12,13^ These procedures, however, become less efficient with increasing system size. A more sophisticated treatment uses SAXS data during sampling. This idea has been applied to MD simulations schemes, either in the form in guided sampling ^14^ or aiming at force-field refinement.^15^ By assembling fragments based on the predicted secondary structure (similar to the ones we use in the present work) and calling CRYSOL in every sampling step, Gajda et al. ^16^ managed to generate good predictions for structures of up to 70 nucleotides. Similarly, Dzananovic et al.^17^ applied such an approach to a viral 3-way junction of roughly 50 nts length. Their tool, called RNA Masonry, also performs fragment assembly but uses the SimRNA^18^ energy function in addition to CRYSOL for the evaluation of every structure. For larger RNA molecules, the program RS3D^19^ can be used, employing a coarse-grained (CG) representation. In the same spirit, the CG model HiRe-RNA has also been extended to guide its simulations with an energy function dependent on SAXS intensities on structures up to 77 nucleotides. ^20^

Here, we present an approach that is suitable for large RNA structures due to the use of a coarse-grained representation of the fragments based on the secondary structure, which is hierarchically refined going through a higher resolution coarse-grained model until atomistic resolution. We incorporate SAXS data via the pair distance distribution function at every sampling step, which is more intuitively accessible to non-crystallographers than the intensity distribution in reciprocal space.

In addition to the possibility to search for a single best structure, we also describe an ensemble-based approach, which optimizes the SAXS profile of the ensemble as opposed to optimizing an individual structure. This way we account for the intrinsic flexibility of large RNA molecules in solution. In the refinement step, we propose a method for removing clashes and fixing broken bonds, and pay special attention to the entanglements, both in their detection and removal, and in the possibility of forming tertiary contacts.

The pipeline starts from the secondary structure and SAXS data, which is used for defining a helix-based representation in an improved version of the ERNWIN model^21^ for exploring and adjusting the global shape of the RNA molecule. Afterwards, the best results are refined by means of the SPlit and conQueR (SPQR) model,^22^ a nucleotide-level resolution coarse-grained description. Finally, the structures are backmapped into an all-atom representation for further MD simulation in explicit solvent. This multiscale approach integrates the strengths of each representation in a way that it resembles the hierarchical folding of RNA.

## MATERIALS AND METHODS

In this section we will first briefly describe ERNWIN, and describe the new features we have added to it and explain the incorporation of SAXS data into its energy function. Later, after a brief introduction of the SPQR model, its capabilities used into the refinement procedure will be exposed. Finally, we present the way our pipeline is assembled and tested.

### Coarse grained structure prediction by ERNWIN

Our in-house RNA structure prediction tool ERNWIN^21^ assembles RNA 3D structures from fragments of known crystal structures, which are defined via the secondary structure elements (stems, interior loops, etc). A rigid coarse-grained representation of these elements is used during sampling.

As described previously,^21^ stems are parametrized by their length and the orientation of the minor groove along the helix. By assuming that each base pair of the helix contributes equally to the helix rotation, we can calculate “virtual” residue and atom positions, which are in good agreement with the true positions. These virtual residue positions can be calculated on the fly and do not have to be stored. More interesting for sampling than stems are the internal loops which connect the stems and introduce angles and offsets between them. Interior and multiloops are parametrized by the distance between and relative orientation of the adjacent stems as well as the relative location of the stems’ minor grooves.

#### Ernwin energy terms

We have previously shown how the reference ratio method can be applied to the sampling of RNA structures: ^21^ For a given measure of interest, such as the radius of gyration, we supply a target distribution of values taken from solved RNA structures. During sampling we compare the distribution of this measure in our sampled structures to this target distribution and calculate a pseudo-energy based on their difference. We use this pseudo-energy in a Metropolis-Rosenbluth-Hastings like accept-reject step and in this way fit the distribution of the measure of interest over the sampled ensemble to the target distribution, irrespective of the distribution this measure would have without the use of an energy.

During the last years, we have retrained the energy potentials on the representative set of RNA 3D structures. ^23^ Furthermore, we have adapted the contributions of long range interactions (loop-loop interaction and A-Minor interaction) to the energy function to better suit longer and more extended RNA molecules. See Supplementary Data Section 1.1 for more details.

For fitting SAXS data, we replaced the radius of gyration energy by a pair distance distribution energy. See section *Fitting the ensemble to the SAXS derived PDD* below for details.

#### Handling of multiloops

A 3-way junction has three single-stranded regions, but is fully defined after sampling only two single-strand fragments, as there are no degrees of freedom left for the third fragment. In such situations, ERNWIN originally only restricted the length in 3D space that is spanned by the last segment (which we will call *“broken” multiloop segment*) and did not perform any sampling. Unfortunately, this approach led to a bias in the distribution of sampled multiloop topologies and generation of unrealistic conformations (see Results).

Here, we explore new ways of assigning a fragment to the *broken multiloop segment* and accept or reject the multiloop conformation based on its fit. We choose to calculate this fit based on the deviation between the first stem of the multiloop and the way we would place this exact same stem after going around the junction according to the sampled fragments. See the Supplementary Data Section 1.2 and Figure S1.1 for details on how we assign fragments to the *broken multiloop segment* and how we benchmark them.

#### Move sets

The most straightforward way to go from one structure to the next during sampling is by changing a single fragment. Due to the fragment based approach, even a single fragment change can potentially change completely the overall conformation, e.g. when we change the angle between two arms of an RNA. These big steps help with quickly exploring the conformational space. However, as the individual parts of the RNA molecule are not independent, in particular if they are linked in a multiloop, big changes to certain fragments have a high chance of introducing clashes or breaking the multiloop constraints, especially in large molecules with complex secondary structure. As ERNWIN rejects structures with clashes, this type of sampling is often inefficient.

For the present version of ERNWIN we have added and tested 5 alternative move types, which are all based on the idea of moving more than one fragment at a time. Most suitable for simple junctions is to move 2 or 3 connected fragments of the same multiloop at a time. For very complex structures we implemented a move type that consists of an exchange of a single fragment followed by a relaxation step based on the clash and multiloop constraints. The other move types and the way we benchmark them are described in the Supplementary Data Section 1.3.

#### Fitting the ensemble to the SAXS derived PDD

The observed SAXS pattern in reciprocal space can be converted to the pair distance distribution function (PDD) in real space via a Fourier transform, and creating this pair distance distribution function is a standard part of SAXS analysis using tools like GNOM.^24^

We interpret the PDD as a distribution of distances between the atomic centers, which we approximate by using 1 or 3 points per residue (depending on the length of the RNA).

We implemented 2 energy functions which act on the PDD in different ways: Either as a potential on the area between the two PDD-curves, or by applying the reference ratio method to each histogram bin of the PDD individually. The latter approach is more powerful, as it operates on the ensemble of sampled structures instead of just a single structure. See the Supplementary Data Section 1.4 and Figure S1.2 for implementation details of these approaches.

### Nucleotide-level refinement

We employ the SPQR model to refine the reconstructed decoys at nucleotide level. The model represents each nucleotide by a phosphate bead and an anisotropic particle which represents the nucleoside. The interactions allow to form stacking, canonical and non-canonical base pairs, and base-phosphate pairs, while the sugar pucker and glycosidic bond angle states are additional degrees of freedom of each nucleotide,^22^ and can be allowed to change dynamically along a simulation.

In the current framework, the structures with broken bonds and clashes, which have an undefined SPQR energy, can be treated with special energy terms which pushes the involved particles towards a finite-energy region. The definition of the energy terms and parameters are described in the Supplementary Data Section 2.1. This relaxation is, however, not enough to guarantee the aptness of the model, since it is common that during the assembly steps the secondary structure elements become entangled. SPQR detects and attempts to remove these artifacts, and later proceeds to perform a search for the possible tertiary contacts proposed by Ernwin. These procedures are described below in more detail.

#### Link detection and removal

To detect and fix entangled secondary structure elements such as hairpins, stems and internal loops, we represent each of them as a closed ring, as depicted in Figure 1a)-e). In practice, these rings are constituted by the segments that join the positions of consecutive phosphate, sugar beads and closing base pairs. We then apply two approaches, previously reported in the literature,^25,26^ which are combined for efficiency and robustness.

**Figure 1:**
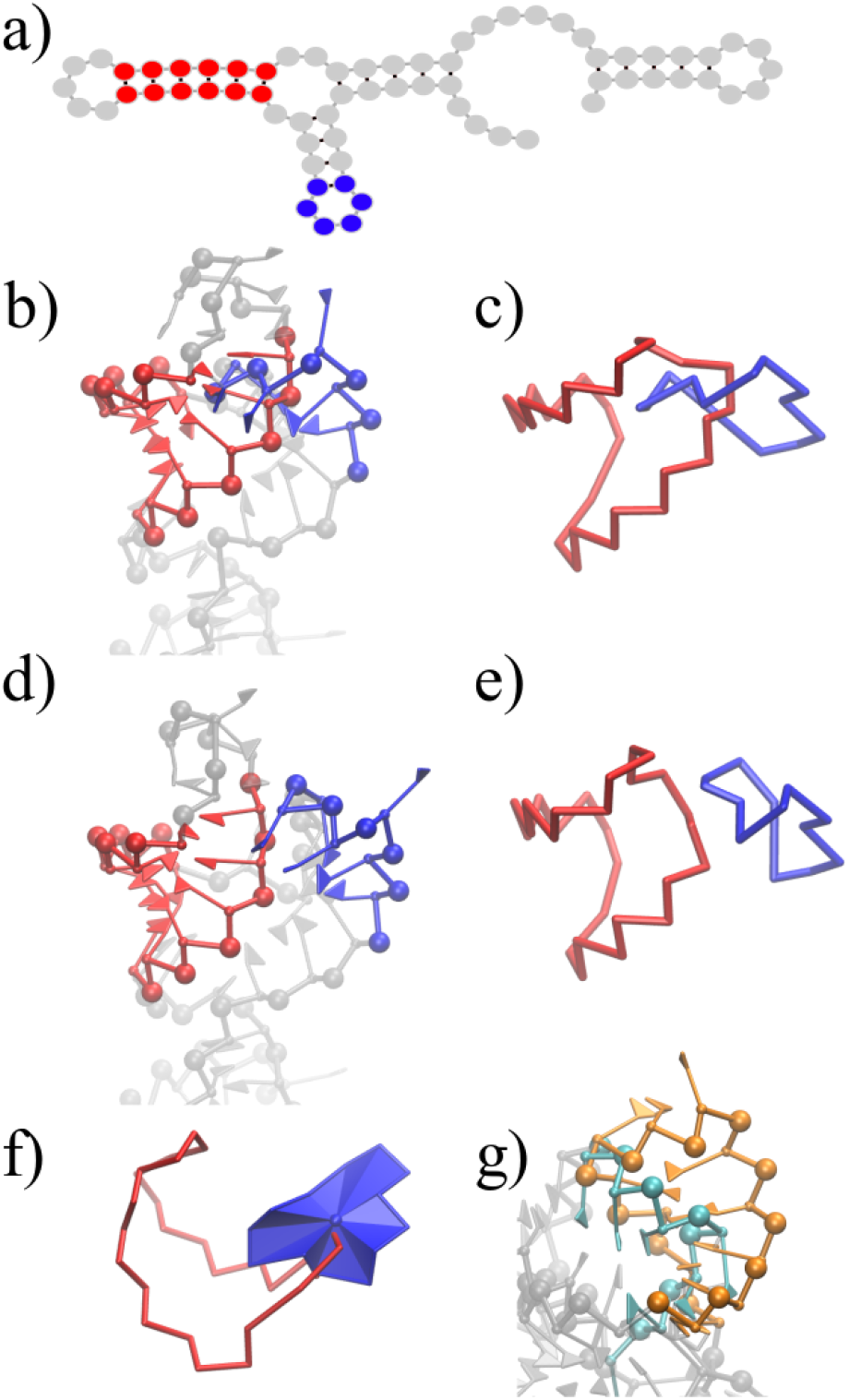
Examples of links for identification and removal a) Secondary structure, b) a hairpin and a stem linked, c) ring representation of the link, d) after clash and link removal and e) ring representation. f) Top view of the link, with triangulated surface of hairpin and g) interpenetrated hairpin and stem, not detected with Gauss integral method.

The first approach makes use of a known way in knot theory of checking whether two rings are linked by evaluated the linking number *L*, defined by the Gauss linking integral:

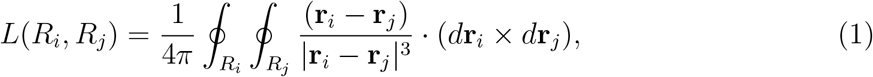

where the vectors **r**_*i*_ and **r**_*j*_ go along the rings *R*_*i*_ and *R*_*j*_ respectively. *L* is the number of times that one ring winds around the other one, which is zero if the rings are not linked. In our case, we evaluate the integrals numerically along segments defining our ring elements. This approach has also been used in the study of the topology of lasso-type proteins with disulfide bonds^25^ and to characterize the topology of proteins.^27^

The second approach, which has also been tested in proteins^26^ and nucleic acids,^28^ consists in evaluating the number of times that the segments which compose a ring pierce the surface enclosed by another ring. In the present case, we consider such a surface as a set of adjacent triangles, formed by two consecutive vertices and the center of mass of the ring; a similar but simpler implementation than the one reported in. ^26^ Such a representation is illustrated in Fig. 1f).

The Gauss integral procedure is in general slower, but it allows to include more complex objects in the analysis, such as three- and four-way junctions, which can also be defined as closed rings and can be difficult to define as a set of triangles due to their geometry. The piercing method, on the other hand, permits the detection of rings crossed by elements such as internal loops which can not be included into a closed ring or short backbone segments between consecutive stems. Most of the occurrences detected by this approach correspond exactly to the links detected by the Gauss integral evaluation, while false positives usually stand for the interpenetration of hairpins and stems of spurious origin, which often involve a large number of clashes in an intricate manner as shown in Fig. 1g). After the links are detected, a repulsive energy term is imposed between the nucleotides involved and specific virtual sites defined on each ring over a short simulation, which guides the relaxation process towards a disentangled conformation. An illustration of the different types of links mentioned before as well as the definition of the virtual sites and repulsive energy terms for each kind of loop are contained in Supplementary Data Section 2.2 in detail. In the current pipeline, the piercing method is used to detect the links, while both piercing and Gauss methods are employed after the relaxation process to confirm the aptness of the model.

#### Tertiary contact search

SPQR allows the introduction of a harmonic potential energy on the ERMSD between the simulated system and a reference structure (see Supplementary Data Section 2.3). This energy term can also be applied on arbitrary sets of nucleotides, which have been used for for enforcing secondary structure elements ^29^ or complex geometries as in intraviral RNA^30^. A short annealing simulation explores the conformational space around the unlinked, refined structure, in search for base pairs between nucleotides belonging to secondary structure elements which interact according to the ERNWIN energy function. The geometry of the found contacts is enforced locally by a hard ERMSD restraint,^31^ while the rest of the nucleotides are pushed towards the original structure with a softer restraint. Parameters are described in Supplementary Data Section 2.4. The result is a structure without clashes nor links; with the fulfillment the possible tertiary contacts at a nucleotide resolution, and with a global structure as close as possible to the one proposed by ERNWIN.

Finally, an atomistic model is obtained from the assembly of template nucleotides on each of the coarse-grained sites, taking into consideration the glycosidic bond angle and sugar pucker conformations. This step usually introduces minor artifacts such a broken covalent bonds, which can easily removed by a short all-atom MD relaxation. The structure can be further refined by introducing an explicit solvent, which is explained in detail in Supplementary Data Section 2.5. The resulting structure is thus suitable for further calculations of SAXS spectra including solvent effects.

### Benchmark against solved PDB structures

To benchmark our pipeline against PDB structures from the protein database, we used the reference structure’s exact coarse grained pair distance distribution function (PDD) for the ERNWIN potential and the reference structure’s secondary structure as basis for the RNA model. We chose the following 4 structures for our benchmark: 3R4F (66 nts), 4PQV (68 nts), 2R8S (120 nts) and 1L9A (126 nts) based on their length and the richness of their secondary structure. The selected structures do not have pseudo knots or large domains interacting with proteins. We then simulated 15 trajectories for each PDB structure with an energy function consisting of 3 components: The pair distance distribution energy, the A-Minor energy and the Loop-Loop interaction energy with different weights for each trajectory. As successful sampling with a strict separation between fragment library and benchmark structures have been shown previously^21^ and we are now more interested the SAXS fitting and sampling efficiency, we now used a full fragment library from the whole representative set of RNA structures. We sampled 20000 to 25000 steps for each structure, which is more than should be necessary for small to medium sized RNA molecules. Simulations took between 12 and 36 hours with one core used per trajectory.

We then selected two structure sets from each trajectory for refinement with SPQR: the structure with the lowest RMSD (to see the best structure Ernwin can produce, to test the ability of ERNWIN to explore the conformational space) and the structure with the lowest pair distance distribution energy.

### Benchmark against real SAXS data

We searched the SASBDB^32^ for SAXS data of RNA molecules which would work well to benchmark our pipeline. Luckily, at the time this paper was started, a recent addition to this database, SASDK34, was ideal for this task: it contains a reasonably sized RNA molecule (118 nt) which is free of G-quadruplexes and overly complicated pseudoknots, and the corresponding paper^33^ contained a predicted all-atom structure which we could use for comparison with our results.

For our benchmark, we performed simulations starting from two different secondary structures: One with only the NMR based secondary structure restraints from Supplementary Figure 7 of the paper we compare our results to (Soni *et al*. ^33^) and one with additional base pairs added by RNAfold^34^ on top of the NMR based restraints.

We performed simulations using ERNWIN with an energy contribution from the experimental pair distance distribution using 3 points per nucleotide in the ERNWIN model. For each secondary structure, 32 simulations were performed, half of which using simulated annealing and half with a constant energy. Simulations used different weights for the energy function. The simulation lengths were set to 1500, 3000 or 5000 steps. We reconstructed a full-atom structure every 25 steps and calculated the CRYSOL 3.0 *χ*^2^ for all of the all-atom structures. Stacking was assigned to the helices by manual inspection of predicted structures.

Later, we used SPQR to refine the last and lowest energy structures of each trajectory of the set of structures with additional RNAfold base pairs.

### Evaluation of *χ*^2^

All reported *χ*^2^ values were calculated by CRYSOL 3.0^35^ over all-atom structures, which can be generated by the Ernwin reconstruction or a backmapping after SPQR refinement.

## RESULTS

### Improvements to the Ernwin method

We could improve the efficiency of sampling and the quality of the predicted structures compared to previous versions of ERNWIN: By replacing distance-only junction constraints with fragment based multiloop constraints, we strongly reduce the bias in the sampled junction topologies. This can be seen from the more structured (less uniform) distribution of angles between adjacent stems, even if they are only connected by the *broken multiloop segment*, as shown in Figures S3.1 and S3.2 and Supplementary Data Section 3.1.

Additionally, we could improve the sampling efficiency and reduce the risk of getting trapped in a local minimum, by introducing new move types which change more than one fragment at a time. We can see that with the new move types, more unique multiloop conformations can be sampled in the same computation time (see Tables S1 and S2 and Supplementary Data Section 3.2 for a direct comparison).

### Benchmark of our pipeline against published PDB structures

The benchmark of our pipeline against real PDB structures showed two main results:

1. For the majority of structures, at least some trajectories converged towards a conformation with a low RMSD and good fit to the native structure.
2. We also had at least one trajectory for each PDB structure that did not converge towards the global minimum but to an alternative conformation. These conformations often have a different arrangement of helices which lead to a similar overall shape. This explains why they are hardly distinguishable from the native conformation based on the pair distance distribution function alone. We will discuss this aspect in detail below.

These two results make it clear again that ERNWIN is a great tool for exploring the conformational space, but additional filtering of the results is needed to find the correct structure. When we use real SAXS data, we use CRYSOL for this filtering step. Table 1 shows the characterizes the sampling trajectories in terms of RMSD to the native structure. Structure 1L9A is an example of a structure where Ernwin performs very well: It has a length of 126 nts, is highly aspherical, making the SAXS profile and pair distance distribution function of the structure very distinctive, and it has a single multiloop with only 3 arms which could potentially clash. The median and average RMSD after the last step of the Ernwin simulation were 11.9 and 15.4 Å, which is very good considering the length of the structure. The best scoring (lowest PDD energy) structure of all trajectories had an RMSD of 7Å. These results show how well Ernwin really can perform under good conditions. Figure 2 panel **a** shows a plot of the RMSD against the sampling step for all individual trajectories, whereas panel **b** shows one sampling trajectory for this structure in more detail.

**Table 1:** RMSDs of predicted structures to the native PDB structure. The first column shows the RMSD of the structure with the best PDD, while the second column shows the best RMSD in the whole ensemble. The last three columns describe the candidate set, which is constructed by taking the best PDD structure of each trajectory. By using one structure per trajectory, we guarantee that they are completely independent of each other. Here we give the best, average and worst RMSD in this candidate set. This shows that by constructing such a candidate set, we usually have at least one structure with an RMSD similar to the best overall RMSD in this small pool of candidate structures.

| PDB | best PDD | best RMSD | min | avg | max |
| --- | --- | --- | --- | --- | --- |
|  | structure<br>(Å) | structure<br>(Å) |  |  |  |
| 1L9A | 8.299 | 5.41 | 7.24 | 16.29 | 36.35 |
| 2R8S | 15.782 | 15.198 | 15.78 | 25.80 | 32.12 |
| 3R4F | 7.439 | 3.975 | 5.87 | 9.08 | 13.77 |
| 4PQV | 13.391 | 8.387 | 8.39 | 12.75 | 17.26 |

**Figure 2:**
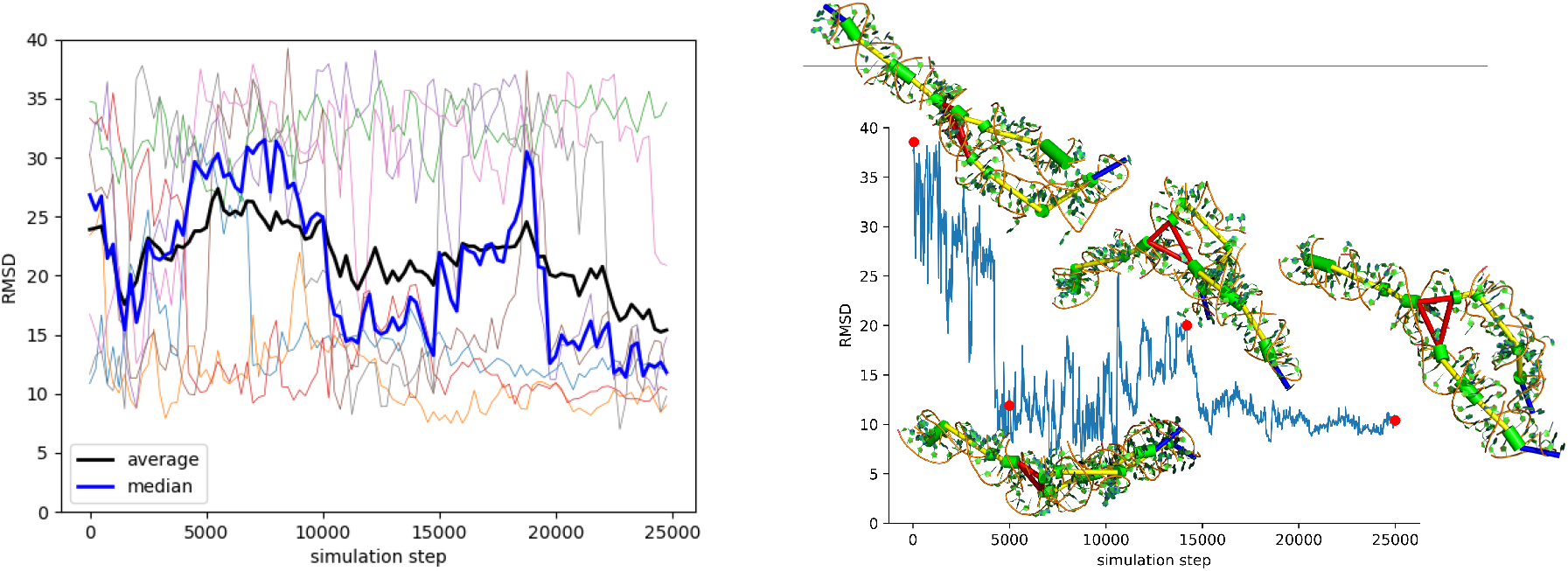
**a** The RMSD over the course of 8 trajectories, plus the median and average RMSD. **b** A successful trajectory, with some of the sampled conformations.

Due to its length and secondary structure, 2R8S is the most challenging of the 4 structures we took for our benchmark. The predicted structure with the best PDD energy has an RMSD of 15.78 Å, which is also very close to the best RMSD overall found in the ensemble. This structure shows a good overall agreement of the global shape and conformation, but noticeable differences in the positions of individual nucleotides, as can be seen in Figure S3.3. As with most RNAs, we again experienced some trajectories that got trapped in local minima with a reasonable PDD energy but bad RMSD and overall shape. For the characteristic interior loop that introduces a 180 degrees turn (*i1*), we have 43 fragments in our fragment library, and the correct fragment is recovered in 6.6% of all structures over all trajectories and is also present in the best PDD structure.

For 3R4F our benchmark shows lower RMSDs than for the comparable 4PQV structure, because of the distinctive bulge with 5 unpaired nucleotides. For this bulge, only 4 fragments were available in our library, one of which was the correct fragment from 3R4F. This example shows how the knowledge of a single fragment, e.g. via homologies or motif search, could improve the overall predictions, and how ERNWIN could use homologies to its advantage. The predicted structure with the best PDD energy has an RMSD of 7.4 Å to the native structure and is shown in Figure S3.4. One the other hand, 4PQV has no distinctive loops and none of the best PDD structures has fragments from 4PQV used for any interior or multi loop. See Figure S3.5 for an example of a sampled conformation.

SPQR refinement results are summarized in Table 2. A large number of samples present links or piercings which can be detected and fixed by the methods presented here. For 1L9A, there is a structure where a three-way junction is linked with an internal loop, which can be detected but not straightforwardly removed by the means presented here. A minor redefinition of parameters is detailed in Supplementary Data Section 4.1 and Figure S4.1 allows to deal with this situation. 2R8S presents the largest number of links, given its size and compactness. An example of two stems linked is shown in Figure S4.2. From 10 hairpin-stem links detected in the whole pool of structures, 4 of them are involved in ERNWIN contacts, which stresses the importance of the refinement for assessing the reliability of the structure. 3R4F is much less compact, and therefore, less topological artifacts are present, although there are links between two stems connected by an unstructured domain (illustrated in Figure S4.3) and unable to form contacts according to the Ernwin energy function. In 4PQV, the links are frequently found between stems which are connected by an unstructured domain (Figure S4.4). Analysis of the RMSD between the refined and unrefined structures shows that the presence of links does not produce a substantial deformation of the structure. Moreover, the difference in this quantity can be attributed mainly to the clashes and the optimization of flexible loops during the refinement process.

**Table 2:** Analysis of refinement. The set of structures for each PDB has 30 structures. The number of links in the set is denoted by *N*_Gauss_ and *N*_Pierce_, detected using the Gauss integral and piercing methods, respectively. The number of tertiary contacts found in Ernwin is *T*_Ernwin_, while *T*_SPQR_ is this number in SPQR representation after refinement. The average RMSD of structures which were not initially entangled is calculated between the unrefined and refined structures in SPQR format and denoted by RMSD_N_. RMSD_L_ contains only linked structures. Averages are reported with standard deviation.

| PDB | $N_{\text{Gauss}}$ | $N_{\text{Pierce}}$ | $T_{\text{Ernwin}}$ | $T_{\text{SPQR}}$ | $\text{RMSD}_N$<br>(Å) | $\text{RMSD}_L$<br>(Å) |
| --- | --- | --- | --- | --- | --- | --- |
| 1L9A | 7 | 12 | 95 | 30 | $3.3 \pm 1.4$ | $3.2 \pm 0.9$ |
| 2R8S | 15 | 23 | 128 | 69 | $4.2 \pm 1.5$ | $4.3 \pm 1.1$ |
| 3R4F | 2 | 7 | 0 | 0 | $1.8 \pm 0.1$ | $2 \pm 0.2$ |
| 4PQV | 5 | 16 | 13 | 5 | $2.2 \pm 0.8$ | $2.5 \pm 0.8$ |

The number of contacts predicted by Ernwin and found in the exploration of SPQR is between 32% and 48%. Nevertheless, in 1L9A one of the A-minors present in the native structure is successfully recovered in 3 instances of the 6 where Ernwin predicts it (see Figure S4.5), while for 2R8S, only one native A-minor is recovered in one Ernwin structure, which is also observed in the SPQR refinement, shown in Figure S4.6.

Further contact analysis of the structures shows that the contact search greatly improves when the sugar pucker and glycosidic bond angle states are allowed to change. In fact, from a set of 150 structures selected randomly from the trajectories of 2R8S and 4PQV, the fixation of the pucker and glycosidic bond angle reduces the number of tertiary contacts found from 76 to 56 and from 183 to 97, respectively.

Finally, we see from Table 2 that the refinement procedure does not introduce a large deviation in the global structure. Visual inspection suggests that the highest RMSD differences correspond to the arrangement of clashed loops in general. Moreover, when introducing the all-atom details, the average clashscore decreases from 115.5 to 5.84 for the four structures described here (details are presented in Table S4 in the Supplementary Data). This opens the way for a further analysis of the SAXS data with a full description of the solvent using more sophisticated approaches which are beyond the scope of the present work.

### Benchmark of our pipeline against real SAXS data

We also benchmark our pipeline against real SAXS data for the *plasmodium falceparum* signal recognition particle ALU RNA, which was deposited in the SASBDB database under the id SASDK34. Soni *et al*. ^33^ predicted ab initio structures using the FARFAR webserver, selected the best cluster of structures based on scoring with CRYSOL (*χ*^2^ = 4.18) and then fitted the structures to the experimental SAXS data using SREFLEX^36^ reaching a *χ*^2^ of 2.0 for their best model.

In our benchmark 13 out of 32 trajectories for the RNAfold secondary structure reached a *χ*^2^ below 2 at least somewhere in the simulation and 4 trajectories had a *χ*^2^ below 2 at the end of the simulation. We refined the best and the last structure of each simulation with SPQR.

For all 13 structures with an initial *χ*^2^ below 2, the removal of clashes and links increased the *χ*^2^, but for roughly half of them the subsequent MD run reduced it again. At the end, 4 out of the original 13 structures still had a *χ*^2^ below 2, two of which had a value below 1.7 (see Figure 3 for an example). The best structure could be further improved using SREFLEX ^36^ to a *χ*^2^ of 1.378.

**Figure 3:**
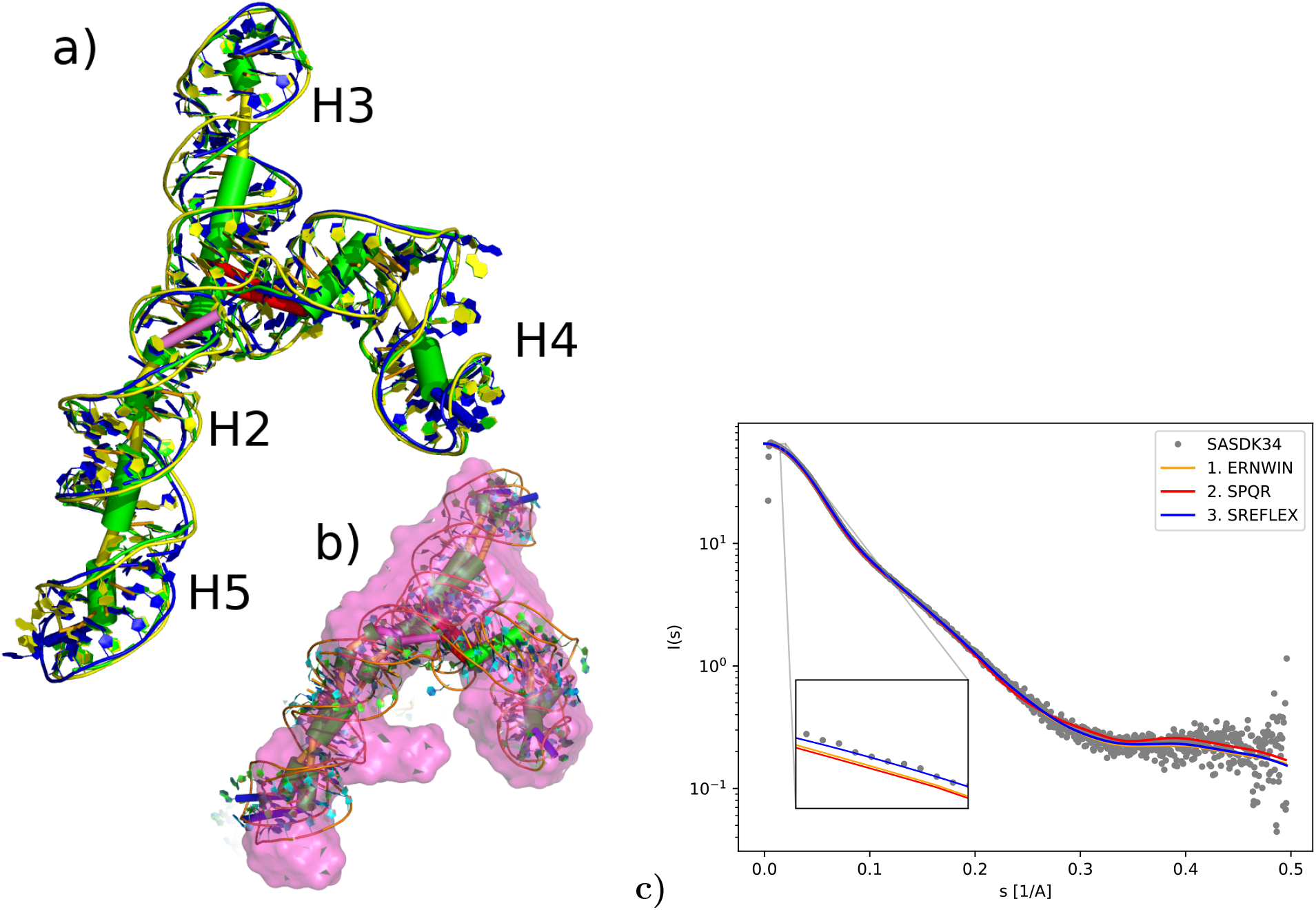
**a)** Structure with the best *χ*^2^ for SASDK34. Blue is the original ERNWIN prediction, green the structure after refinement and yellow the final structure after MD. **b)** The two independent predictions with final *χ*^2^ below 1.7 show a similar conformation with an RMSD of 8Å to each other. In the image, they are both aligned to the volume calculated by Soni *et al*. ^33^ (not to each other) **c** The predicted scattering curve for the best ERNWIN prediction plotted in front of the experimental data from Soni *et al*. ^33^. At small values of *s*, the plot for the unrefined ERNWIN and refined SPQR structure are almost identical, while the SREFLEX curve is slightly closer to the experimental data (see inset), whereas for larger values of *s* the original ERNWIN and the SREFLEX curves are very similar, and the intermediate structure after SPQR refinement deviates a bit from them.

Interestingly, the conformation of our best structure is different to the best prediction reported by Soni *et al*. ^33^. While they predict a stacking between helices H3 and H4, we find a better fit for stacking of helices H3 and H2/H5, but still confirm the overall Y-like shape. Considering the near-equal length of the 3 arms and the fact that we are operating in a *χ*^2^ range that indicates a “not bad fit” but does not indicate “the correct model”, it is not surprising that multiple different conformations and arrangements of the 3 arms can lead to good fits. Additionally, one must not forget the intrinsic flexibility of RNA, especially in such open conformations. The recorded SAXS spectrum might arise from a dynamical ensemble, not a single structure. Thus such SAXS based all-atom predictions have to be combined with biological insights and additional experiments to find the correct solution.

Out of the 13 best trajectories which had at least 1 structure with a *χ*^2^ below 2 before refinement, 4 clearly and 2 somewhat show stacking between H3 and H2/H5 whereas 3 show a stacking of H4 and H2/H5 (with *χ*^2^ of 1.442, 1.459 and 1.996). For 3 structures the local stacking at the multiloop does not correspond to the overall helix orientation and for 1 structure no stacking could be assigned. Our best prediction with an architecture similar to the prediction of Soni *et al*. ^33^ has a *χ*^2^ of 3.0, and could be refined with SPR and SREFLEX to a *χ*^2^ of 1.89. After refinement with SPQR and MD, 4 structures still have a *χ*^2^ below 2, three of which show stacking between H3 and H2/H5, while for one structure the local stacking did not correspond to the overall helix orientation.

The links found and removed with SPQR are less frequent in this case, given the lack of compactness. The few cases observed involved adjacent elements or elements connected by a short unstructured fragment, which did not exhibit tertiary interactions. From the pool of 64 structures, only 5 had links detected with the Gauss integral method and 15 with the piercing method. The average RMSD between the refined and unrefined structures was of 2.4 ± 1 Å and 2.2 ± 1.1Å, for the linked and not-linked models. The secondary structure without additional RNAfold basepairs overall shows similar results, but only 4 trajectories reached a *χ*^2^ below 2. As ERNWIN (in contrast to other tertiary structure prediction tools) does not introduce additional base pairs, it is expected that the RNAfold secondary structure would lead to better predictions.

Finally, we used ARES^37^ to score our best prediction as well as several predicted alternative conformations with low *χ*^2^ to assess the reliability of our models by other methods. This tool returned a predicted RMSD of 8.46 Å (after SREFLEX) for our best prediction and 8.48Å for our prediction that closest matches the structure from Soni *et al*. ^33^ (after ERNWIN, with a *χ*^2^ of 3) and values between 6.5 Å and 10 Å for other alternative conformations.

## DISCUSSION

### Handling of junctions and ergodicity

Junction and pseudo-knot conformations are crucial for the overall conformation of the RNA molecule. In a system with fragments based on secondary structure elements, sampling junctions is quite challenging. For this reason, the authors in Laing *et al*. ^38^,^39^ have developed a random forest approach that classifies the junction family and assigns a geometry before sampling the rest of the structure with their tool RAGTOP.^40^ In contrast to this, it has always been an important feature of ERNWIN to give junctions the flexibility to change during sampling. Here we successfully integrated this into the framework of a fragment based approach by assigning fragments to all junction segments.

The sampling of multiloops as individual segments in ERNWIN requires the introduction of a constraint energy. Together with the clash energy, this causes certain conformations to have an infinite energy and therefore to be forbidden. This means that it is no longer clear whether the sampling is ergodic, as the forbidden regions of the conformational space might for some RNA secondary structures separate the conformational space into disjoint regions. We overcome this limitation by using new move types which exchange more than one fragment at a time, thus opening new paths around the forbidden zones of the conformational space.

### Quality of the pair distance distribution energy for fitting towards SAXS data

The benchmarks have shown that the use of the PDD energy enables the model to sample structures in good agreement with experiment. Nevertheless we observe that robust predictions require one to perform multiple simulation runs to make sure the correct structure is among those that were generated. In addition, we recall that multiple structures might have degenerate PDD-energy, thus making it difficult to identify the correct one.

This situation is especially likely to happen for structures with several arms of nearly equal length, which exhibit a certain symmetry at the level of helix representation. For example, the structure 1Y26 (which was used exclusively in the parametrization of ERNWIN) represents a junction with 3 arms of similar length. Figure 4 shows the native structure, a sampled structure with low PDD energy, and low RMSD and a sampled structure which has a low PDD energy despite having a high RMSD. This structure is a good example of structures where the loop-loop interaction energy of ernwin helps favoring the correct structure more than the PDD energy does.

**Figure 4:**
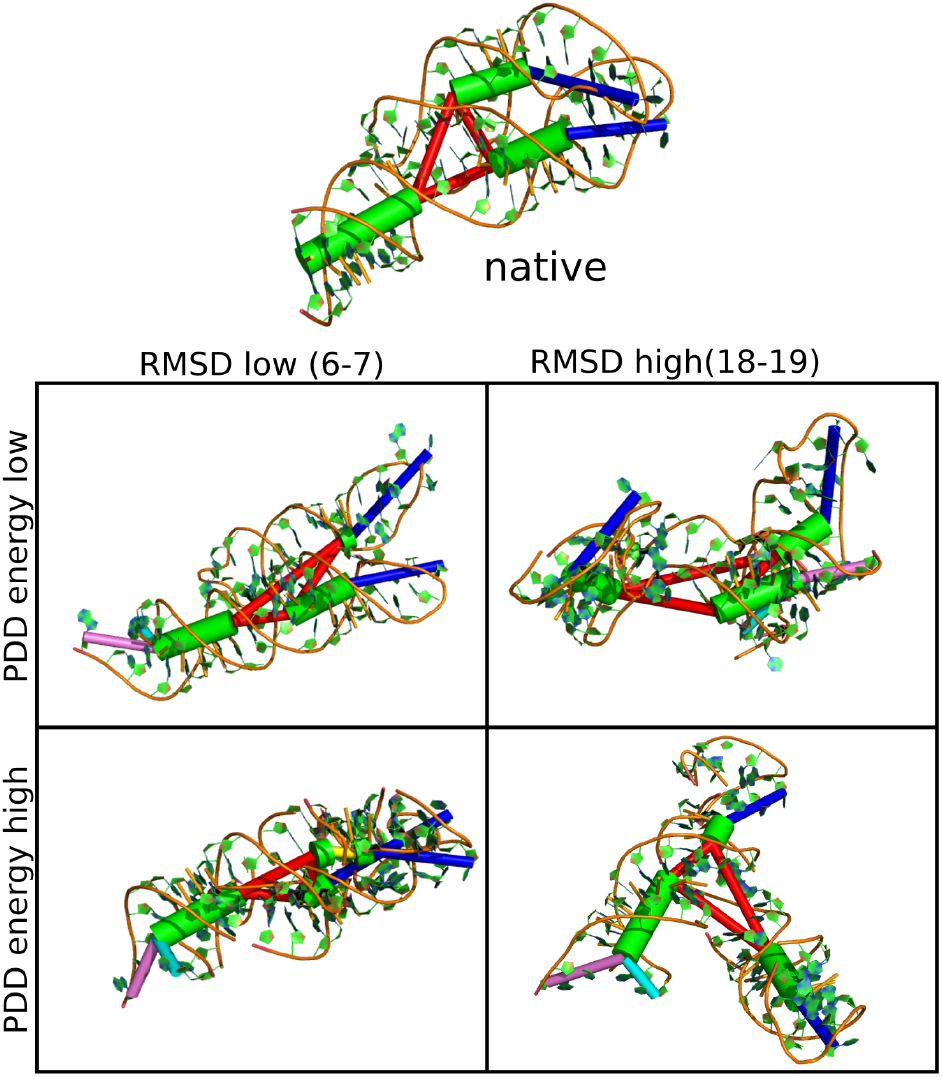
Sampled structures for **1Y26**: It can be seen that the structure at the top right of the table (low PDD-energy, high RMSD) adopts a similar global shape as the native structure, but the arms of the junction are swapped. While the two hairpins are kissing in the native structure, they are at opposing ends in this high-RMSD structure. On the other hand, the structure at the lower left (low RMSD, high PDD energy) looks overall similar to the native structure, but is more compact than the native structure. Note that this figure shows structures before refinement with SPQR.

Still, these limitations of the energy function made it necessary to include a filtering step based on the *χ*^2^ score into our pipeline. On the other hand, the energy is good enough to reliably generate faithful structures in a significant percentage of the trajectories, thanks to the ability of ERNWIN to quickly and efficiently explore the conformational space.

### Why report a unique structure

Although ERNWIN is able to use ensemble based energies, we have chosen to present a single “best” structure for two reasons: First of all, individual structures as illustrative examples of the ensemble are commonly used in biology to simplify things. For instance, this is commonly done when reporting minimum-free-energy secondary structures for RNA. Secondly, it is hard to validate ensemble predictions. On the other hand, individual structures can be easily compared to high resolution crystal structures.

### SAXS and SPQR

We intentionally do not constrain the SPQR refinement with SAXS data. ERNWIN alone does a good job at predicting structures which fit the experimental SAXS pattern well, because SAXS measures a global effect which does not depend on the local orientations of individual nucleotides, which will be refined with SPQR. On the other hand, SPQR does not greatly distort the structures and can turn some models into a more realistic fashion. We see that the good *χ*^2^ of our best prediction is retained after the refinement, while for some other predictions of lower quality the refinement destroys the agreement with the SAXS data. It would be interesting to see if this step of refinement and relaxation without SAXS constraints helps to filter out false positive predictions of ERNWIN and to avoid overfitting, but this will be the subject of future investigations due to the lack of experimental structures with both SAXS and 3D structural data available at the moment.

## CONCLUSION

We introduced here a truly multiscale methodology to propose and reconstruct large RNA structures subject to SAXS experimental restraints. Our approach shows how the conformational space can be explored efficiently using the assembly of secondary structure based fragments, which has been improved for sampling realistic multiloop conformations and for enhancing the exploration of the conformational space by using different types of moves.

By refining the structures with SPQR, it is possible to remove artifacts induced by the coarseness of ERNWIN, which deal with the clashes, topology and the possibility of implementing the tertiary contacts at a nucleotide-level resolution. Moreover, this refinement can be extended to other fragment assembly methods, which are also affected by similar artifacts.^26^ The pipeline ends with the structures refined in all-atom representation in explicit solvent, which allows to analyze further effects of the buffer into the SAXS profile.^10^

Overall, our multiscale method takes the best of each representation, and constitutes a powerful tool in the structure prediction problem of RNA macromolecules in solution.

## ACKNOWLEDGEMENTS

S. P. thanks Prof. Judit Lisoni for infrastructure support and the project FONDECYT Iniciación en Investigación No. 11181334 for financial support.

