## Supplementary Data for "Sampling globally and locally correct RNA 3D structures using ERNWIN, SPQR and experimental SAXS data"

#### 1 Supplementary methods - Ernwin

##### 1.1 Ernwin energy function: Loop-loop interaction energy

In the original `ERNWIN` publication, we used the reference ratio method to fit the distance between hairpins like loop-helix orientations to a target distribution. While this worked well for RNA molecules up to 200 nucleotides, it caused problems with longer molecules, where it disallowed isolated hairpins (with a distance above 100 Å to the next hairpin). With the new versions of `ERNWIN` an alternative energy function is available, which is based on a chemically intuitive binary classification in interacting or non-interacting pairs of loops.

##### 1.2 Handling of multiloops

A multiloop with $n$ single-stranded segments is fully determined after sampling $n-1$ segments and the $n^{th}$ segment does not have to be sampled. Here we introduce a way to still assign a fragment to this *broken multiloop segment*. We choose to measure the fit of the fragment assigned to the broken multiloop segment by the deviation between the stem that would be assigned to it (*virtual stem*) and the way this stem is actually placed. This deviation is measured by the distance between the stems' starts, the angle between them and the difference in twist angle.

Ernwin supports two ways of assigning a fragment to the *broken multiloop segment*:

1. **Searching-Strategy:** Whenever a junction is evaluated, Ernwin can search through all possible parameters for the *broken multiloop segment* and accept the one that minimizes the deviation between the true and the *virtual stem*. (SFJ energy)
2. **Sampling-Strategy**) Or Ernwin can treat the *broken multiloop segment* like any other segment and sample a single fragment for it. (FJC energy)

In both cases the constraints on the maximal deviation between the stem and the virtual stem are then evaluated (implemented a junction constraint energy) and structures with a too high deviation are rejected. Obviously the second strategy (Sampling-Strategy) makes rejects by far more likely than the first.

We benchmark the different approaches by sampling a 3-way junction with 3 equally long multiloop segments and equally long stems. Ideally, all 3 multiloop segments should be indistinguishable and have the same distribution of conformations. The only thing that breaks this symmetry in our model is the fact that we have one *broken multiloop segment*, and this introduces a bias into such loops. We measure this asymmetry using the distribution of angles between adjacent stems as a 1-dimensional proxy for the distribution of multiloop segment conformations. As in our model and for the given scenario, the artificial bias introduced by the broken multiloop segemnt is the only thing that makes these distributions distinguisable, we know that more similar distributions for all 3 segments mean better constraint energies.

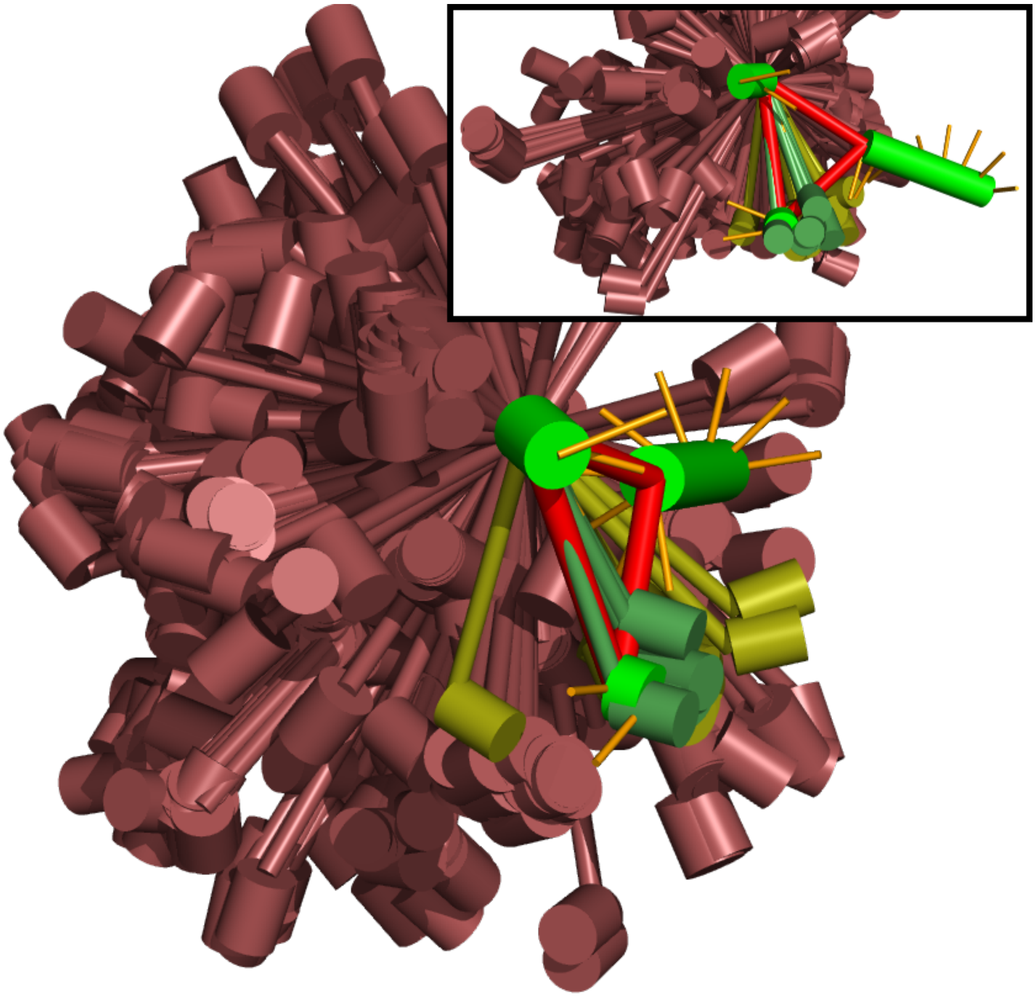

Figure S1.1: Fragment based multiloop constraint: The native conformation of the 3-way junction is shown in bright colors (green for stems, red for multiloop segments). Additionally, possible assigned multiloop segments and their corresponding virtual stem are shown in dimmer colors: green if they deviate at most by 8 Å and 32 degrees, yellow if the deviation is at most 12 Å and 48 degrees and red otherwise. The inset shows the same model from a different angle.

#### 1.3 Move sets

For ERNWIN, we have implemented and benchmarked the following movers in addition to the **SimpleMover** which just changes a random fragment:

The **NElementMover** and the **OneOrMoreElementMover** change two or more random elements at once before evaluating the constraints and energy functions. The **ConnectedElementMover** is a version thereof that changes $n$ elements that are connected to each other. The **WholeMLMover** is very similar, but instead of using a fixed number of fragments to change, it moves all fragments of the multiloop at once.

The **EnergeticJunctionMover** changes 2 or more adjacent single stranded regions of a multiloop. In a single Monte Carlo step it tries all combinations of fragments for the selected single stranded regions until the multiloop fulfills the constraint. This allows for computational optimizations compared to doing a full Monte Carlo step for every combination of fragments. With this move type, version 2 of the new multiloop constraint (assigning a single fragment to the *broken multiloop segment*) becomes somewhat feasible.

The **MoveAndRelaxer** not only addresses the difficulty to fulfill the junction constraints but also the challenges in fulfilling the clash constraints, which mostly come up for larger structures. After sampling a single fragment, we try to change the remaining fragments as needed to remove clashes and fix junction constraints. This is done in a way inspired by the (full) cyclic coordinate descent [3, 1] algorithm and can be seen as a kind of gradient walk. We note that after removal of the *broken multiloop segments*, the Bulge Graph[7] is a rooted tree, where the root node is the element that is first placed (typically the stem "s0" placed at the origin). This tree is a spanning tree of the Bulge Graph, which is a minimum spanning tree if we use the lengths (in nts) of elements as edge weight. For each loop that no longer fulfills the constraints due to the change of the first fragment (in case of pseudoknots it can be more than one loop), starting at the smallest loop, we begin relaxation at the segment closest to the minimum spanning tree's root, because changing this segment will have the greatest impact on the broken multiloop segment. For this segment we try fragments until the constraints are fulfilled or we have tried N (currently 75) fragments. In the latter case we pick the best fragment and continue by relaxing the segment next closest to the minimum spanning tree's root. During choosing of the best fragment, removal of clashes between stems *within* the loop we try to close is given priority over minimizing the gap at the *broken multiloop segments*, because the number of changeable segments between the clashing stems (degrees of freedom) is smaller or equal the number of segments in the loop, which can be relaxed to achieve loop closure. After relaxing the junctions, we try to move additional interior loop elements to remove the remaining clashes.

The gradient walk in this order sometimes gets stuck in a local optimum which does not fulfill all constraints, but even for structures with complex pseudoknotted loop structures it finds a valid solution often enough. With this move type, the second type of fragment based constraints is mostly identical to the first type, because the move itself performs searches through the fragments. By choosing a maximum deviation of 12 Å distance between the stems' starts and 48 degrees between the stem angles and twist angle, we typically find a matching combination of fragments in reasonable time, since we can evaluate this property of the junction independent of the rest of the structure.

To benchmark the different move types we used 3 test structures with different complexity: A 3-way junction, chain X of 2GDI, a tRNA (1FIR) as a simple structure with a single 4-way junction and a 3-way junction plus a kissing hairpin pseudoknot (2YGH). We used no energy (and accepted all structures that fulfilled the constraints). As a starting conformation we use the structure with the highest RMSD to the native structure out of 25 random structures. We then let simulations run for 10 hours and counted the number of unique multiloop conformations in the sampled ensemble after 1 minute, 10 minutes, 1 hour and 10 hours. A conformation is considered unique based on the identity of the sampled fragments for all not-broken multiloop segments.

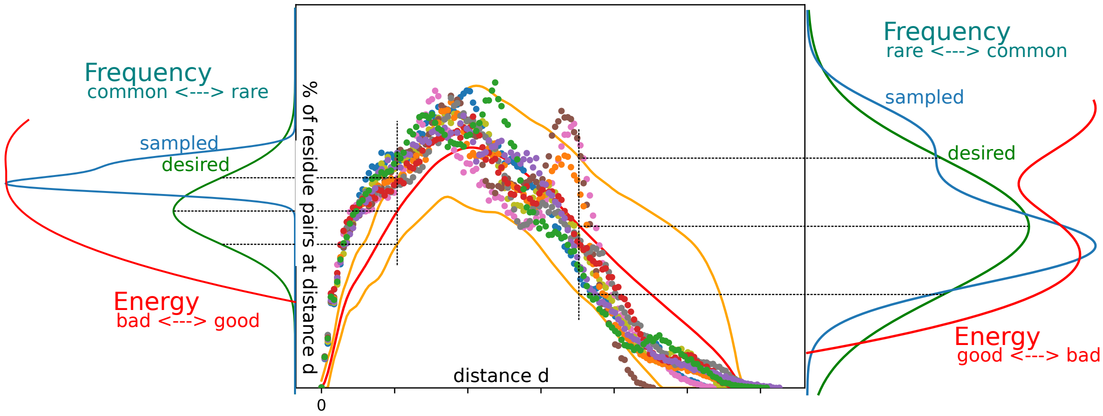


Figure S1.2: **Ensemble based SAXS energy:** The experimental PDD is shown as red line in the central plot. The errors exaggerated by a configurable factor are shown as orange lines. The PDD of the structures that were sampled so far are shown as dots, to indicate that we calculate the PDD in discrete bins. At every sampling step, the energy of the proposed new structure is evaluated in a pointwise fashion for each of the PDD's bins individually. At the left and right side of the central plot, this is indicated for two of those bins. By comparing the distribution of the sampled points to the desired distribution given by the experimental PDD at this distance and the corresponding errors, we can derive an energy function.

#### 1.4 Fitting the ensemble to the SAXS derived PDD

Since the time required to calculate $p(r)$ raises quadratically with the number of points, we approximate the pair distance distribution by fewer points than the number of atoms. Using only the start and end coordinates of stems is insufficient for getting a good approximation of the all atom PDD, so we need more points. For longer RNA molecules it is enough to use 1 point per nucleotide, in our case the the coordinates of the C1' atoms, as we have shown on the example of braveheart[4]. However, we noticed that for shorter RNA molecules, the errors introduced by using only 1 point per nucleotide do not average out as well as they do for longer molecules. Therefor we extended out fragment library to now store 3 point per nucleotide in the local coordinate system of the fragment. While the use of 3 points per nucleotide is common in coarse grained RNA structure prediction, we like to point out that ernwin still uses rigid fragments without inner degrees of freedom and only uses these points for the energy evaluation (calculation of the PDD).

We provide 2 energy functions based on the pair distance distribution. The easier one (called "PDD-energy" simply applies an exponential potential on the deviation between the experimental PDD and the sampled, coarse grained PDD. We calculate this deviation as area between the PDD curves, and since we model the PDD as histogram of distances, this is simply a sum of absolute differences. The second version, the "EPD-energy" (ensemble pair distance distribution) tries to optimize the PDD of the sampled ensemble and not of an individual structure. As our tool especially shines when dealing with larger RNAs, which we expect to be intrinsically flexible to a certain degree, optimizing the pair distance distribution function of the ensemble instead of the PDD of an individual structure better captures reality. We achieve this in the framework of the reference ratio method (as described in the main text) by optimizing the distribution of distributions towards an artificial distribution around the experimental PDD. We do this pointwise by viewing the PDD as a normalized histogram and using a gaussian distribution around the experimental frequency of each bin as target distribution, as illustrated in figure S1.2.

### 2 Supplementary methods - SPQR

#### 2.1 Removal of clashes and reconstruction of broken bonds in SPQR.

The detection of clashes and broken bonds is straightforward in the evaluation of energy of `SPQR`. After a clash detection by the evaluation of the total energy by `SPQR_ENERG`, an energy relaxation is performed by the binary `SPQR_wSA` if there exists any clash or broken bond, which makes use of a particular energy function which tolerates the clashes and helps to remove them. The excluded volume interaction acts on pairs or particles, which can be phosphate, sugars or bases. The sugar particle is in practice a virtual site defined by the position and orientation of the base particle, which is a triangle from where it can be defined a reference frame given its position and orientation. Therefore, the interaction energy between particles $A$ and $B$ is given by

$$U_{exc}(\mathbf{R}_A, \mathbf{\Omega}_A, \mathbf{R}_B, \mathbf{\Omega}_B) = U_{AB}(\mathbf{R}_{AB,A}) + U_{BA}(\mathbf{R}_{BA,B}) \tag{1}$$

where $U_{IJ}(\mathbf{R}_{IJ,I})$ depends on the species of $I$ and $J$, and the vector joining the positions $\mathbf{R}_I - \mathbf{R}_J$ in the reference frame given by the orientation of particle $I$. The functional form of $U_{IJ}$ is

$$U_{IJ}(\mathbf{R}) = \begin{cases} 0 & \text{if } \sum_{ij} a_{ij}\mathbf{R}_i\mathbf{R}_j > 1 \\ \infty & \text{otherwise} \end{cases} \tag{2}$$

where the parameter $a_{ij}$ depend on the species of $I$ and $J$. Note that, for unstructured particles such as the phosphate beads, the above expression is reduced to a hard-sphere potential. In a similar fashion, the potential becomes infinite when a bond between sugar and phosphate beads is broken. A modified version of `SPQR`, included in the simulation package, can deal with these situations by imposing an energy term which does not diverge but pushes particles away when a clash is detected, or favors the formation of a bond when this is broken. In that case, the energy $U_{IJ}$ is given by

$$U^w_{IJ} = C_w(1 - \sum_{ij} a_{ij}\mathbf{R}_i\mathbf{R}_j) \tag{3}$$

where $C_w = 10^5\epsilon_0/\text{Å}^2$, in terms of the `SPQR` energy unit $\epsilon_0$. For bonds and angles, the interaction potentials become

$$U^w_b = C_w(\mathbf{R}_{IJ} - \mathbf{R}^0_{IJ})^2 \tag{4}$$

where $\mathbf{R}^0_{IJ}$ is a reference value for each type of sugar-phosphate bond: intra or inter-nucleotide, and depends on the sugar pucker and glycosidic bond angle state. In the case of angle interactions,

$$U^w_a = C_a(\theta_{IJ} - \theta^0_{IJ})^2 \tag{5}$$

makes use of a reference value $\theta^0$ and a constant $C_a = 10^5\epsilon_0$.

#### 2.2 Link detection and removal in SPQR.

Among the `SPQR` tools, `SPQR_DLINK` is a python script which can detect the links between loops from a given `SPQR` conformation. The format can be easily extended to atomistic PDB structures by means of the script pdb2spqr.py. The script requires the secondary structure in Vienna format from a FASTA sequence, from where the loops are defined in terms of the residue indexes which compose them. These can be hairpin loops (HL), internal loops (IL) and stems (ST). Hairpin loops include also their closing pair, while internal loops are not closed but only a single segment in the piercing method, or closed by considering the stack bounded by the complementary pairs of the first and last nucleotide of the loop.

For the evaluation of the Gauss integral and links in the piercing method, the loops are identified and discretized by the segments joining the consecutive sugar and phosphate beads. In the latter case, each triangle is composed of two consecutive beads and the geometrical center of the loop. The links which have at least one piercing, or a Gauss integral different from zero, are written in an output file which contains the type of each loop and the residue numbers of the nucleotides that constitute it.

The removal is done by a simulation with the `SPQR_wSA` version, which in this case, receives the link input file created by `SPQR_DLINK`. For disentangling the loops, a repulsive interaction is defined between specific virtual sites defined on each loop as follows:

- The geometrical center of loop $i$, denoted by $\mathbf{R}_i$
- The edge site(s) of loop $i$, denoted by $\mathbf{r}_i^j$. For a hairpin loop, there is only one edge site, which is defined as the midpoint between the sugar beads of the closing pair. Stems and internal loops have two edge sites, defined as the midpoint of the closing pairs or by the positions of the sugar beads of its first and last nucleotides, respectively.

The energy function has an additional repulsive energy function between each pair of linked loops defined in the list, by imposing energy terms between virtual sites and modifying the usual energy function. More specifically, the repulsive energy term is exerted between the geometrical center of each loop and the edge sites of its counterpart, as

$$U_{l1,l2} = -K_{ll} \sum_j ((\mathbf{R}_{c,1} - \mathbf{r}_{j,2})^2 + (\mathbf{R}_{c,2} - \mathbf{r}_{j,1})^2. \quad (6)$$

If a stem is involved, an additional repulsive term between the geometrical centers is also added. In addition, the excluded volume interaction between beads of different loops is ommited, which can be written as

$$U_{st1,st2} = -K_{ss} \sum_j ((\mathbf{R}_{c,1} - \mathbf{r}_{j,2})^2 + (\mathbf{R}_{c,2} - \mathbf{r}_{j,1})^2 \quad (7)$$

The energy terms are illustrated in Figure S2.1 for loops found in the samples of the PDB:4PQV.

For linked hairpins, it is clear that a repulsive interaction between the center of mass and closing loops will eventually separate and unlink the loops. The stems are typically more elongated, so they must be pushed away from their center.

The repulsion between the geometrical centers and the closing pairs ensures that the loops will drift away in the right direction. If the edge sites are not included, it is likely that the loops wil dinsentangle towards their side and therefore, the removal of the loop will succeed but creating another link involving a consecutive loop.

The whole procedure is in the script `SPQR_MINI`. Using an atomistic structure as input, it minimizes the energy, removes clashes and links if they are found, to reconstruct the structure in an atomistic pdb.

#### 2.3 $\mathcal{E}$RMSD restrained simulations in SPQR.

`SPQR` has the possibility to apply restraints on the $\mathcal{E}$RMSD, a metric which depends on the relative positions and orientations of the nucleobases which has been shown to be particularly useful for dealing with nucleic acids [2]. In the current version, the restraints can be applied by means of a harmonic potential which depends on this quantity with respect to a reference structure. The structure can be composed of a set of fragments, and therefore, the structures can be enforced locally as in the case of the secondary structure restraints or the tertiary contacts which are composed by three or more nucleotides.

#### 2.4 Tertiary contact search in SPQR.

The search has been performed on the structures with the parameters, and flexibilizing the pucker of the nucleotides which are not involved in the stems. A simulated annealing started from a temperature of $12\epsilon_S$, where $\epsilon_S$ is the `SPQR` energy unit, as described in [6], with 200000 Monte Carlo sweeps for each

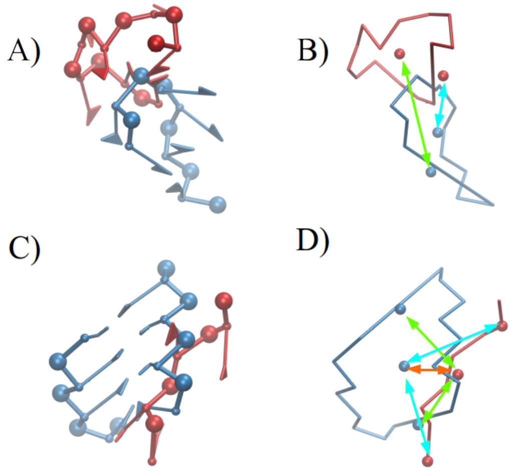


Figure S2.1: Depictions of a structure for link removal: Hairpin-hairpin link in A) `SPQR` representation and B) ring representation, stem-internal loop in C) `SPQR` representation and D) ring representation Ring representations include geometrical centers and edge virtual sites. Note that the repulsive interactions are represented by double arrows in green and cyan for the interaction between a geometrical center and the edge sites of one loop and the other, and viceversa. The interaction between the geometrical centers is colored in orange.

annealing step and a temperature prefactor of $\alpha = 0.75$. The configurations were saved each 10000 sweeps, and all of them were analyzed in search of possible contacts. All the ocurrences of nucleotides corresponding to motifs which interact according to the Ernwin annotations are saved and used as $\mathcal{E}RMSD$ restraints in a later simulation, which enforces these pairs and pulls back the whole structure to its original shape. The simulation is performed at a temperture of $9\epsilon_S$ for 50000 MC sweeps, while the constant $\kappa_{\mathcal{E}}$ has the value of $5000\epsilon_S$ for the tertiary contacts and $5\epsilon_S$ for the global arrangement. As a final step, the energy is minimized by a short Monte Carlo simulation at zero temperature for 10000 sweeps. 10 different random seeds were used in this approach to recover as much contacts as possible, which in some cases, by small perturbations could be missed from the criteria of the `SPQR` annotation tool.

#### 2.5 Backmapping in SPQR.

A brute-force backmapping is finally applied on the final, refined structure, with the largest number of tertiary contacts predicted by Ernwin and no links nor clashes in the `SPQR` representation. In this case, a template atomistic nucleotide is placed on top of each `SPQR` nucleotide, taking into consideration its glycosidic bond angle conformation and sugar pucker. The template is taken from the PDB as a representative nucleotide for each of the states represented in `SPQR`. As an alternative, it can also be taken from the initial structure proposed by Ernwin. The backmapping procedure selects the one that has the lowest RMSD after optimal superposition of the position and orientation of the nucleobase in an `SPQR` representation over the `SPQR` refined structure.

### 3 Supplementary results - Ernwin

#### 3.1 Junction constraints

The distribution of angles between each pair of stems in the 3-way junction with 3 equally long segments should be the same for each pair of stems. However, for the old, distance-based constraints, we noticed that the angles between the stems attached to the *broken multiloop segment* are almost equally distributed, while the angles between the other pairs of stems show a bimodal distribution with one mode corresponding to roughly parallel and one mode to roughly collinear (potentially stacking) orientation.

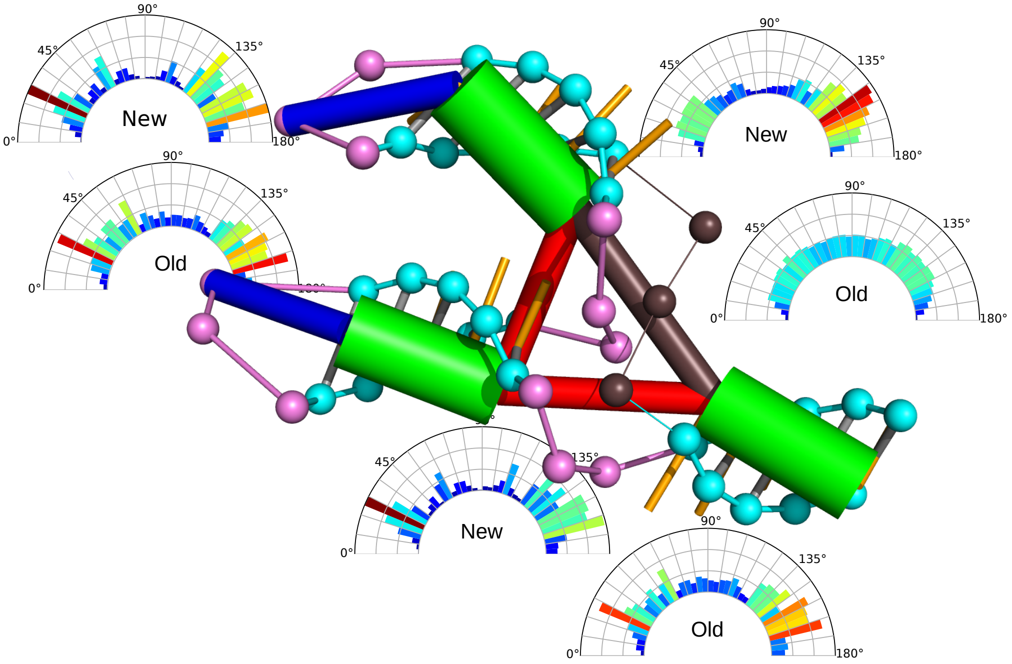


Figure S3.1: Effect of different types of junction constraints. This figure shows an example of a sampled structure of a 3-way junction which for our model exhibits 3-fold symmetry on the level of the secondary structure. The angular histograms describe the sampled distribution of angles between the adjacent stems using two sampling approaches respectively. "New" refers to a fragment-based constraint, while "old" uses only distance-based constraints. It can be seen that the fragment based approach creates the bi-modal distribution of angles (corresponding to a stacking pair of stems) on all 3 junction elements and thus captures the symmetry of the example much better than the old approach.

On the other hand, with the new fragment based approach, we can recover the bi-modal distribution for all 3 pairs of stems. Such a bimodal distribution is typical for experimentally solved structures of 3-way junctions [7, 5] Our new approach only causes a small bias, which manifests in a smoother distribution of angles between the stems connected to the broken multiloop segment. Figure S3.1 shows the angle distribution for both kinds of junction constraints and Figure S3.2 shows a two sample Kolmogorov–Smirnov tests for similarity of the distributions.

It also became apparent that the cutoff of the energy should not be too small, as this would reduce the number acceptable conformations too much and thus impede sampling.

Mover | KS-Test

**NElementMover[3]**

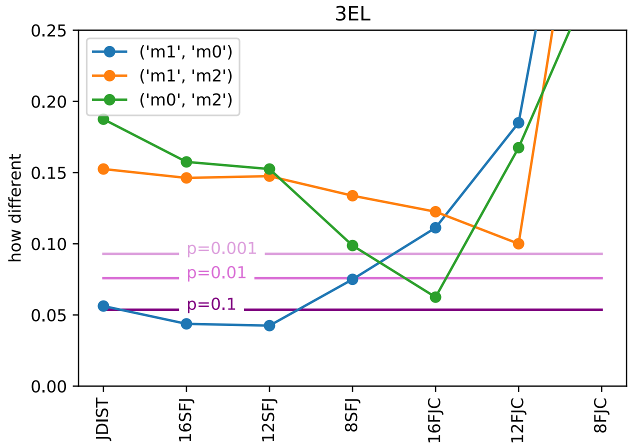


**EnergeticJunctionMover**

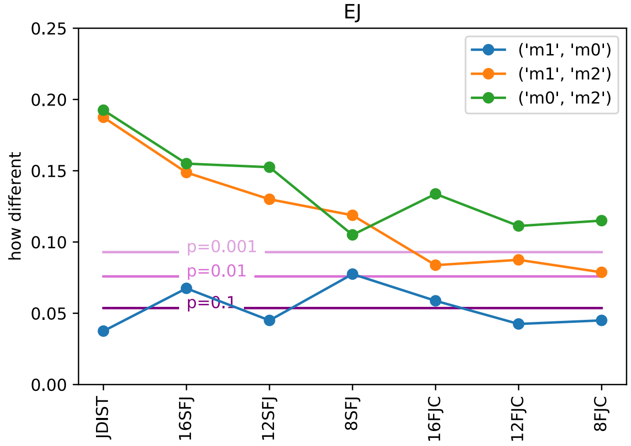


**MoveAndRelaxer**

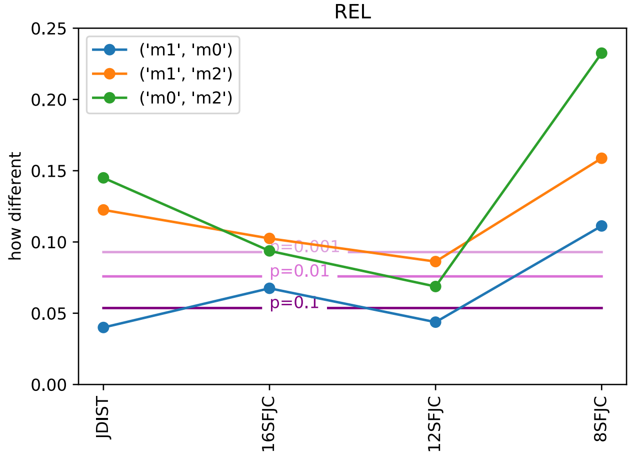


Figure S3.2: Comparison of distributions of angles. For a symmetric 3-way junction, we measure the angle between the stems adjacent to each multiloop segment to produce 3 distributions. Then we compare them in a pairwise manner using the two sample Kolmogorov–Smirnov test. We used different movers (rows in the figure) and different junction constraint energies (x axis of each figure) for sampling and only took every 25th sampling step to achieve independent data points. The left column is the normal KS-test. For the right column the angle distributions were blurred before the KS-test was performed. The junction constraint energies used (= x-labels) were: **JDIST**: distance based constraints, **8FJC**: Fragment based junction closure energy, corresponds to **Sampling-Strategy** at cutoff 8Å and 4*8=32 degrees, **8SFJ**: Searching Fragment based junction closure energy, corresponds to **Searching-Strategy**

#### 3.2 Move types

| | 2GID | | | | 1FIR | | | | 2YGH | | | |
|---|---|---|---|---|---|---|---|---|---|---|---|---|
| | 10 min | | 10 h | | 10 min | | 10 h | | 10 min | | 10 h | |
| | # | % | # | % | # | % | # | % | # | % | # | % |
| Simple Mover | 250 | 1.0 | 16345 | 1.0 | 185 | 1.0 | 17400 | 1.0 | 10 | 1.0 | 1767 | 1.0 |
| 3 Connected Elements | 194 | 0.8 | 39446 | 2.4 | 717 | 3.9 | 43797 | 2.5 | 44 | 4.4 | 2559 | 1.4 |
| Combination | 662 | 2.6 | 46248 | 2.8 | 405 | 2.1 | 28017 | 1.6 | 12 | 1.2 | 6112 | 3.4 |

Table S1: Summary of the benchmark of different move types using a searching based junction constraint. We show the absolute number of unique multiloop conformations (column #) and a comparison to the Simple Mover (column %). See table S2 for a full table including all tested move types and a sampling based junction constraint energy.

The new types of moves for going from one conformation to the next strongly improve the sampling efficiency (See table S1 and S2): Changing 3 connected elements in one move increases the number of unique multiloop conformations sampled in the same time by a factor of 1.4 (2YGH) to 2.5 (1FIR) compared to the simple mover. If we use a combination of movers, including the MoveAndRelaxer, the 3ElementMover and a Simple Mover, among others, we get the best results for more complex molecules. This combination produces 1.6 (1FIR) to 3.4 (2YGH) times more unique multiloop conformations in the same time. The MoveAndRelaxer and combinations of movers using it are most useful for complex multiloops and pseudoknots, whereas the computational time spent on the relaxation step does not pay off that much for simple 3- or 4-way junctions.

#### 3.3 Benchmark against solved PDB structures

### 4 Supplementary Results - SPQR

`SPQR` can remove links between stems, hairpins and internal loops, which can be composed by one or two strands. A junction, which by definition has three or more stems connected by unstructured segments, can be defined as a closed loop and incorporated in the Gauss integral, but an analogous definition of the virtual sites might not be optimal for the removal of the link. In this case, we opted for identifying the single strand which pierces the other loop, and treat it as a link between a single strand and a closed internal loop. This decomposition can be generalized to any type of junction, which will be automatized in future versions of the code. The structure before and after refinement is shown in Figure S4.1.

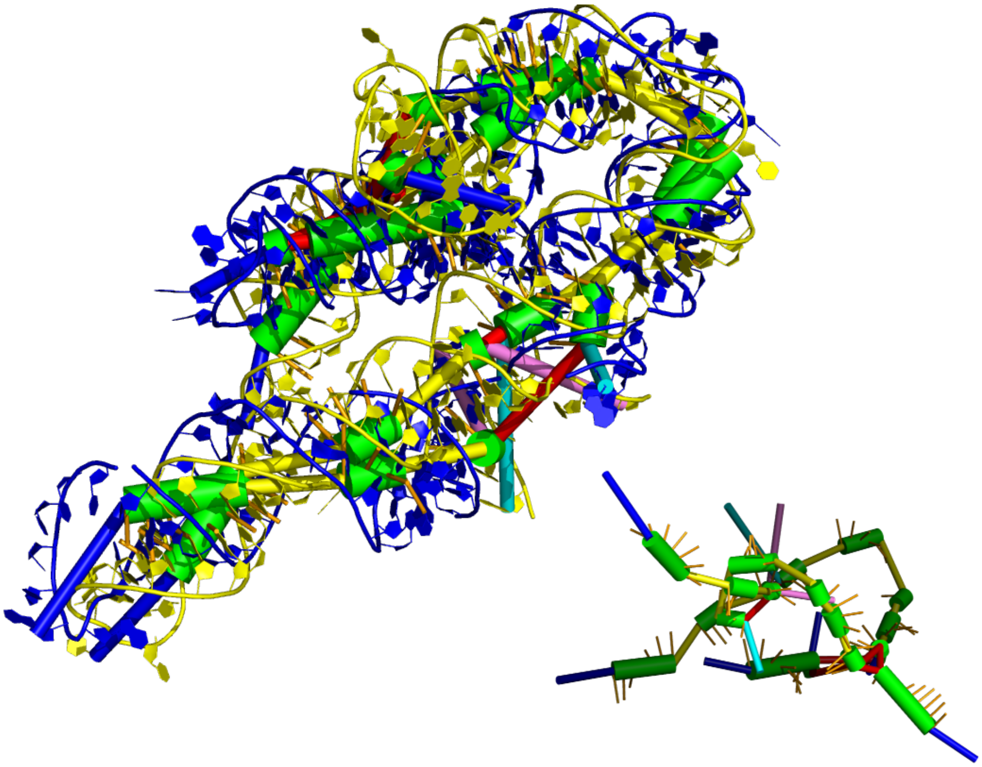

Figure S3.3: **top** Native structure for 2R8S (yellow) and predicted structure with the best PDD energy (blue). **bottom right** A predicted structure with reasonable PDD energy but bad fit (RMSD 32) to the native structure (bright colors) and the native conformation (dim colors).


[5] Aurélie Lescoute and Eric Westhof. Topology of three-way junctions in folded RNAs. *RNA*, 12(1):83–93, jan 2006.

[6] Simón Poblete, Sandro Bottaro, and Giovanni Bussi. A nucleobase-centered coarse-grained representation for structure prediction of rna motifs. *Nucleic Acids Research*, 46(4):1674–1683, 2017.

[7] B. Thiel, I. K. Beckmann, P. Kerpedjiev, and I. L. Hofacker. 3d based on 2d: Calculating helix angles and stacking patterns using forgi 2.0, an RNA python library centered on secondary structure elements. [version 2; peer review: 2 approved]. *F1000Research*, 8:287, April 2019.

| | | 1FIR | | | | | | | |
|---|---|---|---|---|---|---|---|---|---|
| Junction | Move type | 1 min | | 10 min | | 1h | | 10h | |
| FJC | WholeMlMover | 1 | 2% | 1 | 1% | 2 | 0% | 15 | 0% |
| FJC | ConnectedElementMover(2) | 16 | 37% | 73 | 39% | 428 | 26% | 7612 | 44% |
| FJC | EnergeticJunctionMover(2) | 5 | 12% | 29 | 16% | 268 | 16% | 2848 | 16% |
| FJC | SimpleMover | 9 | 21% | 54 | 29% | 279 | 17% | 2925 | 17% |
| FJC | EnergeticJunctionMover(3) | 9 | 21% | 20 | 11% | 77 | 5% | 942 | 5% |
| FJC | EnergeticJunctionMover(whole ML) | 1 | 2% | 13 | 7% | 60 | 4% | 705 | 4% |
| FJC | NElementMover(4) | 9 | 21% | 69 | 37% | 781 | 47% | 11564 | 66% |
| FJC | EnergeticJunctionMover(4) | 3 | 7% | 14 | 8% | 65 | 4% | 670 | 4% |
| FJC | ConnectedElementMover(4) | 8 | 19% | 59 | 32% | 1358 | 81% | 21877 | 126% |
| FJC | NElementMover(3) | 6 | 14% | 23 | 12% | 236 | 14% | 5254 | 30% |
| FJC | NElementMover(2) | 15 | 35% | 64 | 35% | 206 | 12% | 5864 | 34% |
| FJC | Combination of Movers | 6 | 14% | 37 | 20% | 176 | 11% | 1373 | 8% |
| FJC | MoveAndRelaxer | 4 | 9% | 28 | 15% | 191 | 11% | 2215 | 13% |
| FJC | ConnectedElementMover(3) | 13 | 30% | 78 | 42% | 1167 | 70% | 14370 | 83% |
| SFJ | WholeMLMover | 2 | 5% | 20 | 11% | 117 | 7% | 1078 | 6% |
| SFJ | EnergeticJunctionMover(2) | 44 | 102% | 450 | 243% | 2797 | 167% | 27560 | 158% |
| SFJ | EnergeticJunctionMover(4) | 23 | 53% | 305 | 165% | 1840 | 110% | 18894 | 109% |
| SFJ | EnergeticJunctionMover(whole ML) | 36 | 84% | 338 | 183% | 1946 | 116% | 19104 | 110% |
| SFJ | NElementMover(2) | 22 | 51% | 297 | 161% | 2100 | 126% | 24176 | 139% |
| SFJ | ConnectedElementMover(2) | 81 | 188% | 359 | 194% | 2428 | 145% | 24707 | 142% |
| SFJ | NElementMover(3) | 42 | 98% | 619 | 335% | 3331 | 199% | 31393 | 180% |
| SFJ | SimpleMover | 43 | 100% | 185 | 100% | 1671 | 100% | 17400 | 100% |
| SFJ | NElementMover(4) | 66 | 153% | 506 | 274% | 3426 | 205% | 33983 | 195% |
| SFJ | MoveAndRelaxer | 11 | 26% | 135 | 73% | 879 | 53% | 8683 | 50% |
| SFJ | ConnectedElementMover(3) | 109 | 253% | 717 | 388% | 4134 | 247% | 43797 | 252% |
| SFJ | ConnectedElementMover(4) | 77 | 179% | 791 | 428% | 4457 | 267% | 44033 | 253% |
| SFJ | EnergeticJunctionMover(3) | 39 | 91% | 338 | 183% | 2153 | 129% | 21282 | 122% |
| SFJ | Combination of Movers | 41 | 95% | 405 | 219% | 2770 | 166% | 28017 | 161% |
| | | 2GDI | | | | | | | |
| FJC | WholeMlMover | 1 | 5% | 3 | 1% | 4 | 0% | 43 | 0% |
| FJC | ConnectedElementMover(2) | 15 | 79% | 179 | 72% | 1947 | 113% | 13751 | 84% |
| FJC | EnergeticJunctionMover(2) | 25 | 132% | 121 | 48% | 715 | 41% | 6434 | 39% |
| FJC | SimpleMover | 7 | 37% | 119 | 48% | 582 | 34% | 9740 | 60% |
| FJC | EnergeticJunctionMover(3) | 9 | 47% | 70 | 28% | 442 | 26% | 4654 | 28% |
| FJC | EnergeticJunctionMover(whole ML) | 8 | 42% | 79 | 32% | 399 | 23% | 4525 | 28% |
| FJC | NElementMover(4) | 43 | 226% | 318 | 127% | 2044 | 118% | 21810 | 133% |
| FJC | EnergeticJunctionMover(4) | 6 | 32% | 74 | 30% | 471 | 27% | 4789 | 29% |
| FJC | ConnectedElementMover(4) | 16 | 84% | 255 | 102% | 753 | 44% | 12650 | 77% |
| FJC | NElementMover(3) | 14 | 74% | 192 | 77% | 1427 | 82% | 17466 | 107% |
| FJC | NElementMover(2) | 9 | 47% | 188 | 75% | 1377 | 80% | 17566 | 107% |
| FJC | Combination of Movers | 17 | 89% | 183 | 73% | 487 | 28% | 5440 | 33% |
| FJC | MoveAndRelaxer | 9 | 47% | 117 | 47% | 579 | 33% | 6626 | 41% |
| FJC | ConnectedElementMover(3) | 13 | 68% | 357 | 143% | 1525 | 88% | 18909 | 116% |
| SFJ | WholeMLMover | 3 | 16% | 19 | 8% | 178 | 10% | 2701 | 17% |
| SFJ | EnergeticJunctionMover(2) | 17 | 89% | 180 | 72% | 1685 | 97% | 31541 | 193% |
| SFJ | EnergeticJunctionMover(4) | 8 | 42% | 151 | 60% | 1254 | 72% | 27874 | 171% |
| SFJ | EnergeticJunctionMover(whole ML) | 18 | 95% | 148 | 59% | 2329 | 135% | 29076 | 178% |
| SFJ | NElementMover(2) | 29 | 153% | 186 | 74% | 1232 | 71% | 26458 | 162% |
| SFJ | ConnectedElementMover(2) | 18 | 95% | 174 | 70% | 1216 | 70% | 24215 | 148% |
| SFJ | NElementMover(3) | 15 | 79% | 229 | 92% | 1009 | 58% | 35752 | 219% |
| SFJ | SimpleMover | 19 | 100% | 250 | 100% | 1730 | 100% | 16345 | 100% |
| SFJ | NElementMover(4) | 34 | 179% | 333 | 133% | 1364 | 79% | 44093 | 270% |
| SFJ | MoveAndRelaxer | 28 | 147% | 252 | 101% | 1423 | 82% | 13168 | 81% |
| SFJ | ConnectedElementMover(3) | 14 | 74% | 194 | 78% | 3153 | 182% | 39446 | 241% |
| SFJ | ConnectedElementMover(4) | 45 | 237% | 362 | 145% | 3947 | 228% | 43869 | 268% |
| SFJ | EnergeticJunctionMover(3) | 9 | 47% | 190 | 76% | 1834 | 106% | 28436 | 174% |
| SFJ | Combination of Movers | 90 | 474% | 662 | 265% | 4633 | 268% | 46248 | 283% |
| | | 2YGH | | | | | | | |
| FJC | WholeMlMover | 1 | 100% | 4 | 40% | 10 | 10% | 69 | 4% |
| FJC | ConnectedElementMover(2) | 1 | 100% | 4 | 40% | 12 | 12% | 72 | 4% |
| FJC | EnergeticJunctionMover(2) | 1 | 100% | 1 | 10% | 5 | 5% | 79 | 4% |
| FJC | SimpleMover | 1 | 100% | 2 | 20% | 4 | 4% | 102 | 6% |
| FJC | EnergeticJunctionMover(3) | 1 | 100% | 2 | 20% | 3 | 3% | 103 | 6% |
| FJC | EnergeticJunctionMover(whole ML) | 1 | 100% | 2 | 20% | 14 | 14% | 109 | 6% |
| FJC | NElementMover(4) | 1 | 100% | 1 | 10% | 5 | 5% | 114 | 6% |
| FJC | EnergeticJunctionMover(4) | 1 | 100% | 4 | 40% | 19 | 19% | 134 | 8% |
| FJC | ConnectedElementMover(4) | 1 | 100% | 1 | 10% | 7 | 7% | 165 | 9% |
| FJC | NElementMover(3) | 1 | 100% | 3 | 30% | 12 | 12% | 234 | 13% |
| FJC | NElementMover(2) | 1 | 100% | 2 | 20% | 11 | 11% | 408 | 23% |
| FJC | Combination of Movers | 1 | 100% | 2 | 20% | 26 | 26% | 485 | 27% |
| FJC | MoveAndRelaxer | 1 | 100% | 9 | 90% | 53 | 53% | 514 | 29% |
| FJC | ConnectedElementMover(3) | 2 | 200% | 5 | 50% | 7 | 7% | 587 | 33% |
| SFJ | WholeMLMover | 1 | 100% | 4 | 40% | 70 | 70% | 889 | 50% |
| SFJ | EnergeticJunctionMover(2) | 2 | 200% | 4 | 40% | 50 | 50% | 961 | 54% |
| SFJ | EnergeticJunctionMover(4) | 2 | 200% | 10 | 100% | 29 | 29% | 970 | 55% |
| SFJ | EnergeticJunctionMover(whole ML) | 2 | 200% | 9 | 90% | 88 | 88% | 1010 | 57% |
| SFJ | NElementMover(2) | 1 | 100% | 12 | 120% | 148 | 148% | 1344 | 76% |
| SFJ | ConnectedElementMover(2) | 3 | 300% | 31 | 310% | 143 | 143% | 1504 | 85% |
| SFJ | NElementMover(3) | 5 | 500% | 20 | 200% | 173 | 173% | 1590 | 90% |
| SFJ | SimpleMover | 1 | 100% | 10 | 100% | 100 | 100% | 1767 | 100% |

Table S2: Benchmark of different move types, full data. A summary of this data can be found in table S1.

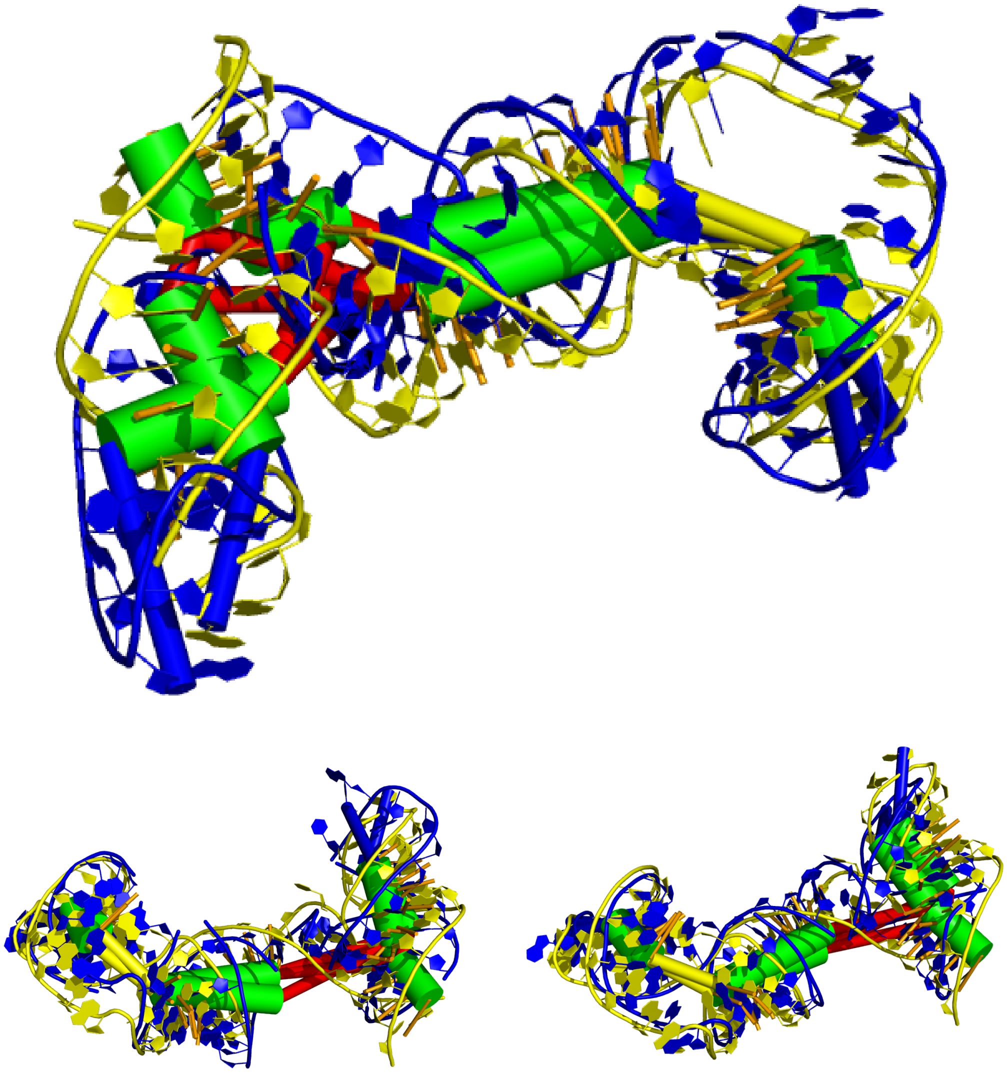

Figure S3.4: Native structure for 3R4F (yellow) and predicted structure (blue). **Top:** The predicted structure with the best PDD energy **Bottom left** The lowest RMSD structure among the set of structures with the best PDD energy of the respective trajectory. In total it had the 3rd best PDD energy value. **Bottom right** The structure with the best RMSD (but with a bad PDD energy)

| PDB | Clash score | | | |
|---|---|---|---|---|
| | Best PDD | | Lowest RMSD | |
| | Unrefined | Refined | Unrefined | refined |
| 1L9A | 127.8 | 7.5 | 124.6 | 6.6 |
| 2R8S | 127.9 | 6.7 | 128.5 | 6.6 |
| 3R4F | 90 | 3.7 | 97.1 | 2.9 |
| 4PQV | 117.6 | 5.8 | 111.6 | 7 |

Table S3: Clash score for the sets of generated structures.

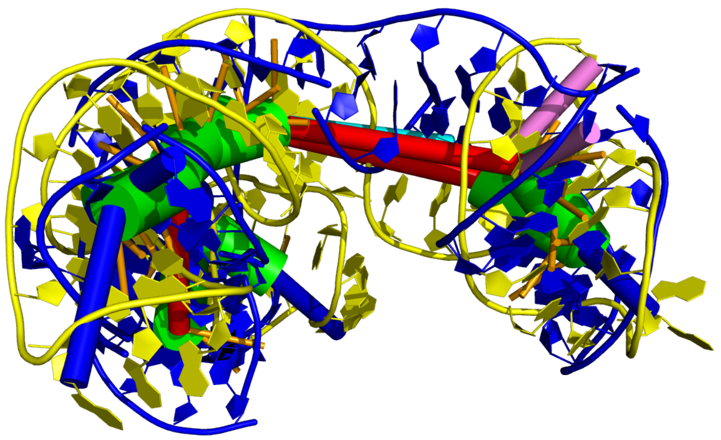

Figure S3.5: Native structure for 4PQR (yellow) and predicted structure (blue) with the best PDD energy.

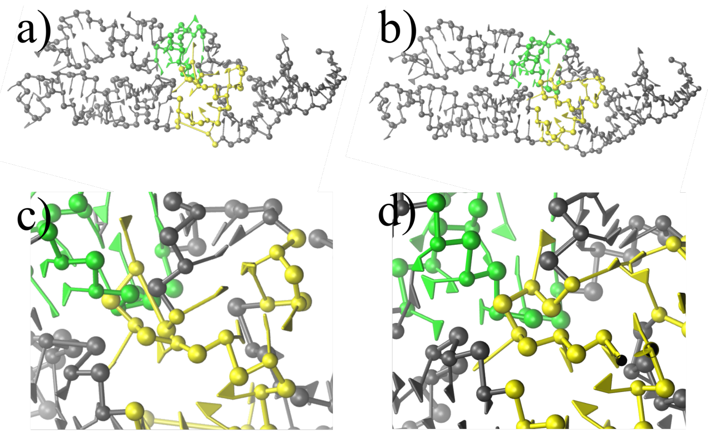


Figure S4.1: For 1L9A: Junction defined by nucleotides 17, 18, 57-66, 109-111 and internal loop defined by 70-76, 101-105: a) Junction-internal loop linked. b) Junction-internal loop unlinked. c) Zoom of a). d) Zoom of b).

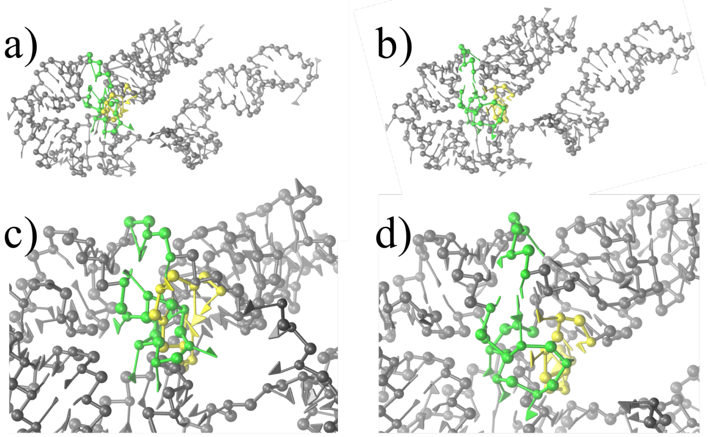


Figure S4.2: For 2R8S: a) Link between s1 and s2. b) $h0$ and $i3$ unlinked. c) Zoom of a). d) Zoom of b).

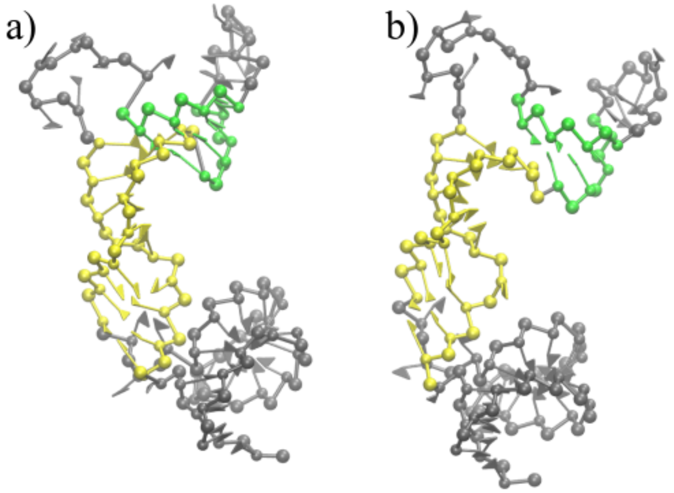


Figure S4.3: For 3R4F: a) Link between $s1$ and $s2$. b) s1 and s2 unlinked.

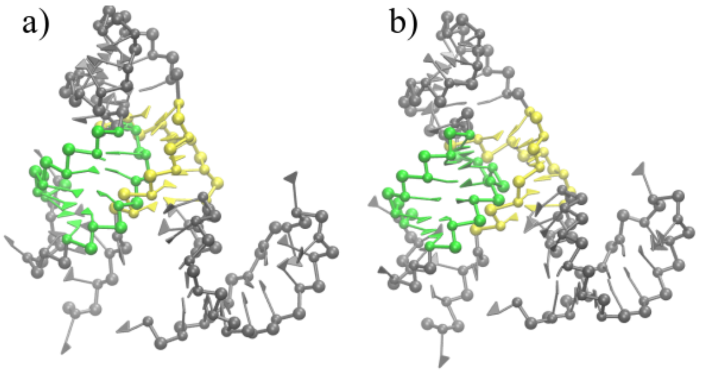


Figure S4.4: For 4PQV: a) Link between $h1$ and $s2$. b) s1 and s2 unlinked.

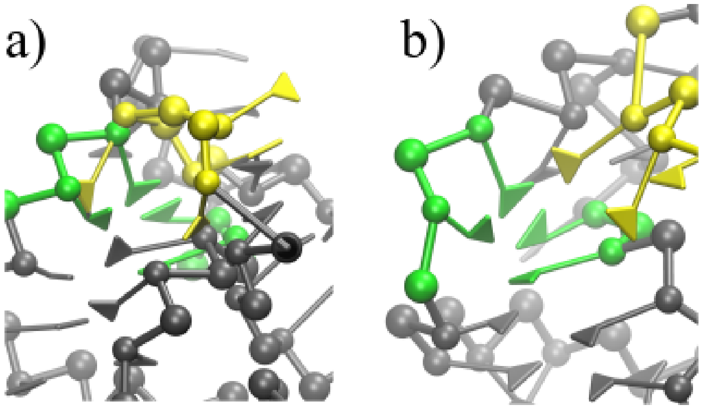


Figure S4.5: Contact between $h0$ and $s9$ in 1L9A a) after Ernwin reconstruction and b) after `SPQR` refinement.

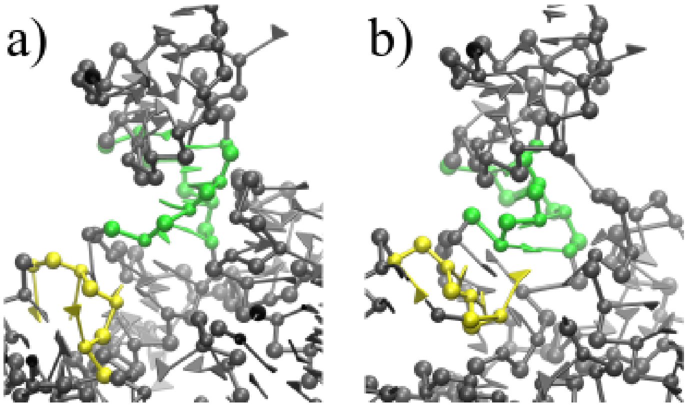


Figure S4.6: Contact between $h0$ and $s8$ in 2R8S a) after Ernwin reconstruction and b) after `SPQR` refinement.
